# Mitochondrial Dysfunction Rewires Macrophage Metabolism, Driving Pro-inflammatory Priming and Immune System Remodeling

**DOI:** 10.1101/2025.11.01.686039

**Authors:** Nikita Markov, Aref Hosseini, Timothée Fettrelet, Kevin Oberson, Anna Brichkina, Shida Yousefi, Carole Bourquin, Hans-Uwe Simon

**Author notes:** **Corresponding authors:** Nikita Markov, Institute of Pharmacology, University of Bern, Inselspital INO-F, CH-3010 Bern, Switzerland., Hans-Uwe Simon, Institute of Pharmacology, University of Bern, Inselspital INO-F, CH-3010 Bern, Switzerland.

## Abstract

Macrophage activation is tightly coupled to cellular metabolism: classically activated pro-inflammatory (M1) macrophages rely on glycolysis and a disrupted tricarboxylic acid cycle, whereas alternatively activated (M2) macrophages depend on oxidative phosphorylation (OXPHOS) and fatty acid oxidation. Although mitochondria are central to this metabolic plasticity, it remains unclear whether mitochondrial dysfunction itself can dictate macrophage polarization.

Using macrophage-specific OPA1 knockout mice, we investigated how mitochondrial dysfunction influences macrophage metabolism and immune homeostasis. Loss of OPA1 caused severe impairment of OXPHOS, reduced mitochondrial membrane potential, and a compensatory glycolytic shift, driving M0 and M2 macrophages toward an M1-like bioenergetic state. Integrative metabolomic and transcriptomic analyses revealed strong priming of OPA1-deficient macrophages towards classical activation, including accumulation of M1-associated metabolites (lactate, succinate, itaconate) and upregulation of NF-κB–driven and other inflammatory gene programs, resulting in increased secretion of IL-6 and TNF even in the absence of stimulation.

Functionally, this metabolic shift primed non-activated and M2 macrophages toward partial M1 polarization with enhanced bactericidal capacity, while simultaneously suppressing M2-associated processes such as proliferation, efferocytosis, and expression of Arg1, CD206, and RELMα.

In vivo, *OPA1^ΔM^* mice displayed reduced peritoneal macrophage abundance, impaired self-renewal after IL-4 complex stimulation, and compensatory monocyte recruitment. The remaining macrophages exhibited increased MHCII and reduced RELMα expression, consistent with partial M1 skewing and loss of alternative activation. These local alterations were mirrored systemically: blood profiling revealed enhanced T-cell activation and sex-specific remodeling of immune composition during inflammation and aging.

Collectively, these findings demonstrate that mitochondrial dysfunction serves as a cell-intrinsic cue that primes macrophages toward a glycolytic, pro-inflammatory phenotype while constraining their M2 properties and proliferative capacities. This dual metabolic and functional rewiring highlights mitochondrial integrity as a pivotal determinant of macrophage immunometabolic identity and reveals how its disruption can reshape both local and systemic immune homeostasis.

## Introduction

Macrophages are immune cells characterized by enormous plasticity and capable of adapting their functional and metabolic programs to the local microenvironment. Depending on the encountered stimuli, they acquire a wide spectrum of activation states broadly referred to as classical (M1) and alternative (M2) polarization(1–3). Classically activated, or M1, macrophages arise in response to microbial products, interleukins and interferons, and mediate pathogen clearance through phagocytosis, production of pro-inflammatory cytokines, reactive intermediates, and tissue-damaging enzymes(4). In contrast, alternatively activated, or M2, macrophages emerge in response to IL-4, IL-13, and other signals, and orchestrate tissue repair, remodeling, and resolution of inflammation(5,6). Although this dichotomy provides a useful conceptual framework, macrophage activation *in vivo* rarely conforms to discrete M1 or M2 identities. Instead, tissue macrophages integrate diverse metabolic and signaling cues to form mixed or transitional phenotypes that are context-dependent and shaped by the metabolic environment, oxygen availability, and systemic processes. Consequently, macrophage polarization *in vitro* often only partially recapitulates their behavior *in vivo*, emphasizing the need for mechanistic models that link metabolic rewiring with functional diversity in physiologic and pathological settings.

A central hallmark of macrophage polarization is its tight coupling to cellular metabolism. Inflammatory M1 macrophages rely predominantly on glycolysis and a truncated tricarboxylic acid (TCA) cycle that supports rapid ATP production and biosynthesis of inflammatory mediators such as succinate, nitric oxide, and itaconate(7–9). Conversely, M2 macrophages depend on oxidative phosphorylation (OXPHOS) fatty acid oxidation (FAO) and glutaminolysis, metabolic programs that sustain long-term biosynthetic, reparative, and anti-inflammatory functions(10–12). The reconfiguration of metabolic fluxes is not merely a downstream consequence of activation but a driving force that determines the transcriptional and functional identity of macrophages. Mitochondria sit at the core of this metabolic plasticity, serving not only as bioenergetic organelles but also as signaling platforms that control calcium currents, generate metabolites and reactive oxygen species (ROS) regulating epigenetics, gene expression, and cytokine production(13–15).

Despite the recognized importance of mitochondria in macrophage activation, it remains unclear how mitochondrial dysfunction itself, independent of classical stimuli, affects macrophage phenotype. On one hand, pharmacological inhibition of the electron transport chain (ETC) or loss of respiratory capacity is known to mimic aspects of M1 activation, promoting glycolysis and upregulating M1-like signaling(16,17). On the other hand, sustained or severe mitochondrial damage has been associated with macrophage exhaustion and impaired immune responses. The question of whether intrinsic mitochondrial defects can actively drive macrophage polarization, and in which direction, therefore remains unresolved.

Two recent studies attempted to address this issue but reached divergent conclusions. Shanshan Cai *et al.* (2023) demonstrated that myeloid-specific deletion of *Ndufs4*, a mitochondrial Complex I component, reprograms macrophages toward glycolysis and amplifies pro-inflammatory cytokine responses upon LPS stimulation(18). In contrast, Ricardo Sánchez-Rodríguez *et al.* (2023) reported that myeloid-specific loss of the mitochondrial fusion protein OPA1 blunts classical activation and enhances the expression of M2 markers, suggesting an opposite, anti-inflammatory outcome(19). These discrepancies underscore the complexity of mitochondrial control over macrophage function and point to the influence of experimental variables such as differentiation state, nutrient composition, and sex. Therefore, it remains unclear whether mitochondrial dysfunction primes macrophages toward a pro-inflammatory or anti-inflammatory trajectory.

In this study, by integrating bioenergetic profiling, metabolomics, transcriptomics, and functional assays in both male and female macrophages, we demonstrate that OPA1 loss in non-activated macrophages (M0) induces profound mitochondrial impairment and metabolic remodeling towards an M1 state, accompanied by NF-κB activation and partial gain of inflammatory phenotype. This intrinsic reprogramming occurs even in the absence of external stimuli and translates *in vivo* into disrupted macrophage renewal, restricted M2 polarization, and altered systemic immune activation. Our findings reconcile previously contradictory reports by showing that mitochondrial dysfunction primes macrophages toward M1 phenotype while simultaneously impairing their M2 functions and proliferative capacities, revealing a dual role for mitochondrial integrity as both a brake and a determinant of macrophage functional identity.

## Results

### Loss of OPA1 induces mitochondrial dysfunction and a glycolytic shift resembling M1 macrophages

Given that mitochondrial dysfunction is closely linked to M1 macrophage polarization, we hypothesized that its induction might predispose macrophages toward an M1-like phenotype. To test this, we sought to induce a comparable level of mitochondrial dysfunction in macrophages without triggering their activation or classical M1 polarization. To achieve a stable, predictable, and reproducible phenotype, we employed a genetic rather than a pharmacological approach. Specifically, we used *Lyz2^Cre/Cre^ Opa1^flox/flox^*mice, in which *Opa1* deletion occurs selectively in macrophages and neutrophils. Loss of OPA1 protein is known to cause profound mitochondrial dysfunction, although the extent of this effect can vary depending on the targeted cell type(20–22). To examine the consequences of OPA1 loss in macrophages, we differentiated bone marrow-derived macrophages (BMDMs) from both male and female *Lyz2^Cre/Cre^* (hereafter referred to as *control*) and *Lyz2^Cre/Cre^ OPA1^flox/flox^*(hereafter referred to as *OPA1^ΔM^*) macrophages. These macrophages were subsequently polarized toward M1 and M2 phenotypes or left untreated (M0) for comparison within each genotype. Bioenergetic profiling using a Seahorse analyser revealed that loss of OPA1 profoundly impairs mitochondrial respiration including both ATP-linked and maximal respiration in macrophages of both sexes under all tested conditions (Figures 1A–C, S1A–C). Notably, in M1 macrophages, OPA1 deficiency further exacerbated the activation-induced impairment of OXPHOS, nearly abolishing mitochondrial respiration. As expected, the reduced energy production resulting from mitochondrial dysfunction was partially compensated by an upregulation of glycolysis in *OPA1^ΔM^* macrophages (Figures 1D–E, S1D–E).

**Figure 1.**
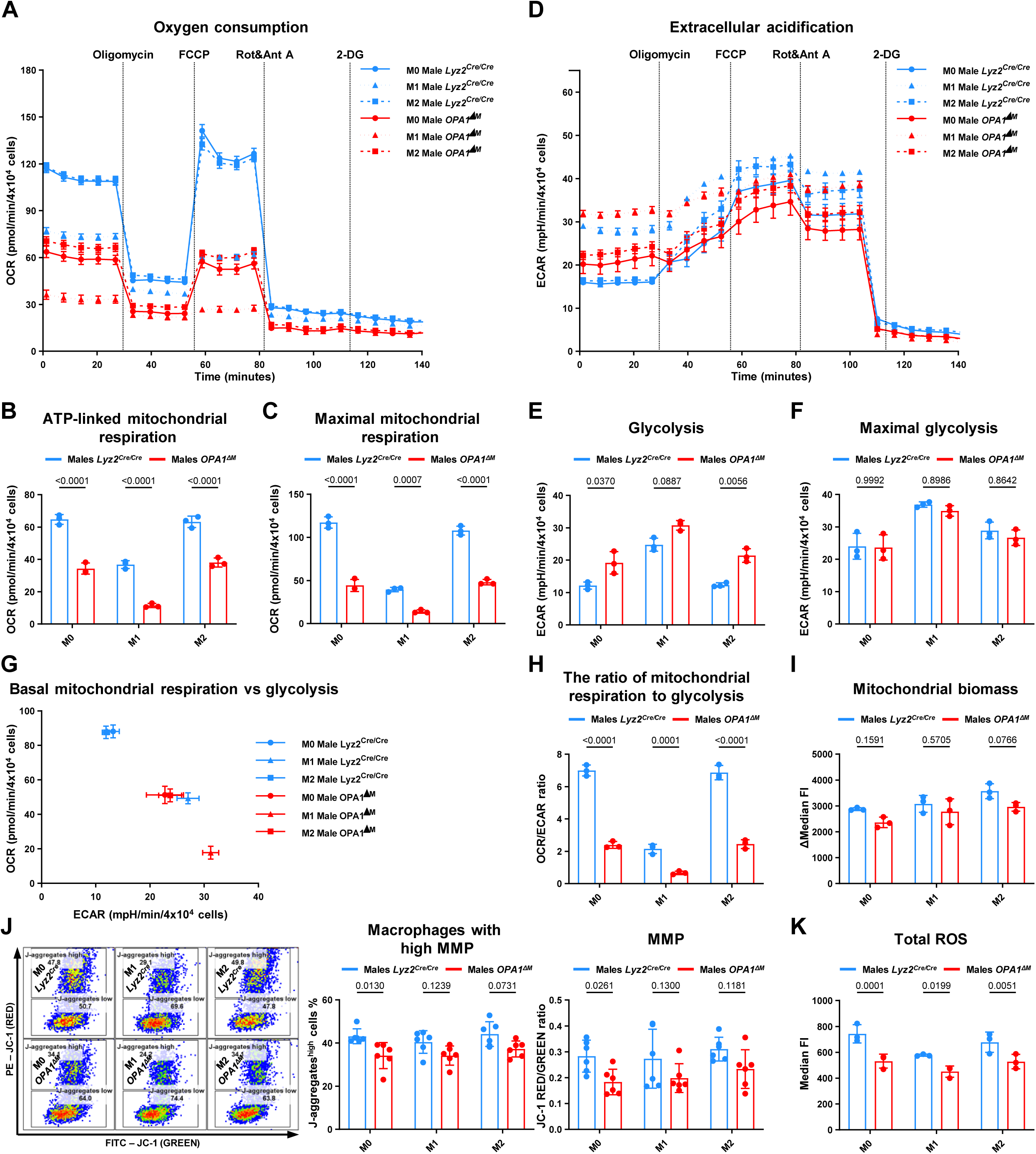
Loss of OPA1 drives a glycolytic shift and reduces mitochondrial respiration in macrophage. (A–K) Control and *OPA1^ΔM^*BMDMs differentiated from *Lyz2^Cre/Cre^* and *Opa1^flox/flox^ Lyz2^Cre/Cre^* **male** mice, respectively, were polarized toward M1 and M2 phenotypes or left untreated (M0) and analyzed after 24 h. (A–H) Bioenergetic profiling using a Seahorse XF analyzer following sequential injections of oligomycin, FCCP, rotenone + antimycin A, and 2-DG. (A–C) Parameters related to oxygen consumption rate (OCR). (D–F) Parameters related to extracellular acidification rate (ECAR). (A, D) Representative Seahorse measurements from one experiment. (B, C, E, F, H) Quantification of OXPHOS- and glycolysis-associated parameters. (I) Mitochondrial biomass determined by flow cytometry using 200 nM MitoTracker Green FM staining. (J) Mitochondrial membrane potential and the proportion of high-MMP cells measured by flow cytometry with 3 μM JC-1 dye. (K) Total ROS quantified by flow cytometry using dihydrorhodamine 123. Data represented as mean ± SD; n = 3–6 (each dot indicates one biological replicate). Statistical significance was determined by two-way ANOVA with Sidak’s correction for multiple comparisons; exact p values are indicated within the panels.

Interestingly, while the maximal glycolytic capacity was comparable between male control and OPA1-deficient macrophages, it was markedly reduced in female knockouts. This observation suggests a potential bottleneck in NAD⁺ regeneration via the ETC, thereby limiting glycolytic flux. Supporting this notion, decreased absolute glycolysis was detected only in female M1 *OPA1^ΔM^* macrophages compared to their control counterparts, but not in M0 or M2 populations.

Although the overall bioenergetic pattern was similar between sexes, female OPA1-deficient macrophages exhibited a less pronounced OXPHOS impairment and a greater reliance on glycolysis, implying that mitochondrial dysfunction and the resulting metabolic adaptation follow distinct dynamics in male and female cells (Figures 1G, S1G). Nonetheless, the metabolic reprogramming induced by OPA1 loss shifted M0/M2 macrophages of both genders towards an M1-like bioenergetic phenotype without immunological activation, as evidenced by a reduced OCR/ECAR ratio reflecting increased glycolytic dependence and suppressed OXPHOS (Figures 1H, S1H).

Interestingly, loss of OPA1 also led to a modest reduction in mitochondrial biomass, most prominently observed in M2 macrophages, which are particularly dependent on OXPHOS. This effect was most pronounced in female M2 cells (Figures 1J, S1J). Measurements of mitochondrial membrane potential (MMP) further revealed a marked decrease upon OPA1 deletion, as indicated by both a reduced JC-1 red/green fluorescence ratio and a lower frequency of cells displaying high MMP. These changes were more pronounced in male macrophages and in subsets relying predominantly on OXPHOS (M0 and M2).

Similarly, mitochondrial ROS, primarily generated by Complexes I and III, were more substantially reduced in male cells, suggesting a sex-dependent effect of OPA1 loss on mitochondrial activity (Figures 1K, S1K). The stronger decrease in ROS levels in male macrophages may indicate that mitochondrial dysfunction modulates redox-sensitive signaling and oxidative stress pathways differently in male and female cells.

Collectively, these findings demonstrate that OPA1 deficiency reprograms macrophage bioenergetics by suppressing OXPHOS and inducing a compensatory glycolytic upregulation reminiscent of M1 macrophages. Notably, the magnitude and metabolic consequences of these changes differ between male and female macrophages, highlighting sex-specific responses to mitochondrial dysfunction.

### Loss of OPA1 metabolically reprograms macrophages towards an M1-like phenotype

Metabolic rewiring in M1 macrophages is tightly linked to their pro-inflammatory functions, as alterations in both metabolic pathway activity and the abundance of effector metabolites critically determine their functional state(11,23,24). To investigate how mitochondrial dysfunction shapes the metabolic landscape of macrophages lacking OPA1, we performed targeted mass spectrometry-based metabolomic profiling of male OPA1-deficient macrophages. After data filtering and quality control, quantitative information was obtained for 173 metabolites representing the substantial fraction of core macrophage metabolome (Figure 2A).

**Figure 2.**
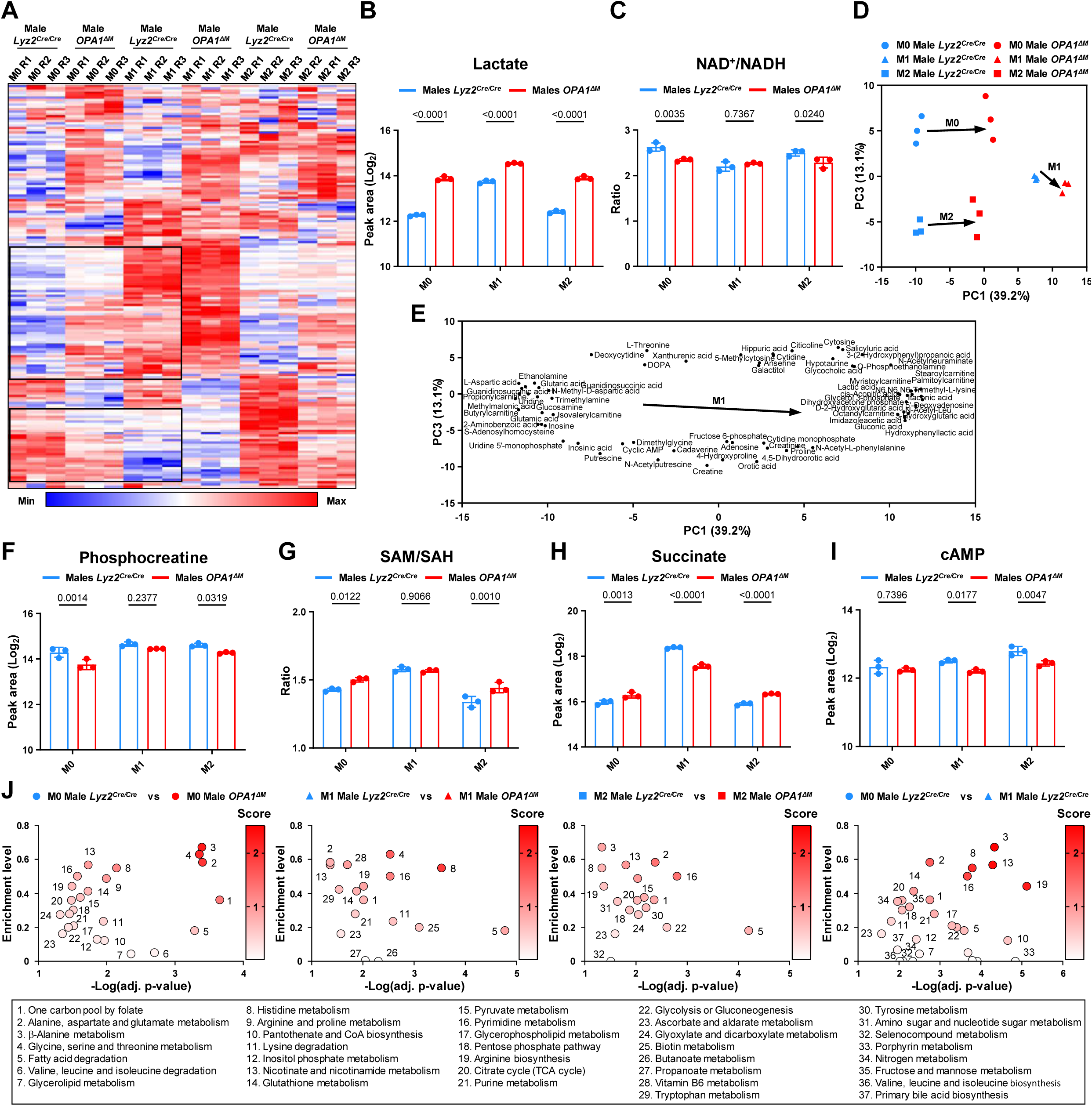
Loss of OPA1 reprograms macrophage metabolism. (A–J) Control and *OPA1^ΔM^* BMDMs differentiated from *Lyz2^Cre/Cre^* and *Opa1^flox/flox^ Lyz2^Cre/Cre^* **male** mice, respectively, were polarized toward M1 and M2 phenotypes or left untreated (M0) and analyzed after 24 h. Targeted metabolomic profiling was performed on cell extracts; 173 metabolites passed quality control and filtering criteria and were included in the analysis. (A) Heatmap illustrating global metabolite changes across genotypes and polarization states. (B, C) Abundance of lactate and the NAD^+^/NADH ratio. (D) Principal component analysis of all samples; arrows indicate the direction of M1-associated metabolic remodeling. (E) The top 10% PCA loadings contributing most to group separation. (F–I) Relative abundances of selected metabolites involved in central metabolic pathways. (J) Pathway enrichment analysis performed using MetaboAnalyst 6.0. Data represent mean ± SD; n = 3 (each dot indicates one biological replicate). Statistical significance was determined by two-way ANOVA with Sidak’s correction for multiple comparisons; exact p values are indicated within the panels.

Given the central role of mitochondria in cellular metabolism, we anticipated widespread metabolic alterations and indeed, OPA1 loss profoundly reshaped the macrophage metabolome. To validate the dataset, we examined metabolites known to reflect shifts in glycolytic and mitochondrial activity, including lactate levels and the NAD⁺/NADH ratio. Consistent with the observed bioenergetic remodeling, OPA1-deficient macrophages displayed elevated lactate accumulation and a reduced NAD⁺/NADH ratio, confirming enhanced glycolytic flux and impaired mitochondrial respiration (Figures 2B–C).

To integrate and compare the metabolic relationships among macrophage polarization states, M1-associated metabolic changes, and those arising from OPA1 deficiency, we performed principal component analysis (PCA) of the macrophage metabolome (Figure 2D). The PCA revealed clear metabolic segregation among M0, M1, and M2 macrophages, reflecting distinct metabolic programs characteristic of each phenotype. Strikingly, both M0 and M2 macrophages lacking OPA1 clustered closer to the control M1 population, indicating that mitochondrial dysfunction drives their metabolomes towards an M1 state. Moreover, *OPA1^ΔM^* M1 macrophages were shifted even further along the M1-associated axis compared with control M1 macrophages, suggesting an exaggerated accumulation of M1-related metabolites. The metabolites contributing most strongly to this separation are shown as the selection of top PCA loadings (Figure 2E).

Although adenosine nucleotides (ATP, ADP, AMP), the most direct indicators of cellular energy status, were not detected in our dataset, we quantified phosphocreatine, which serves as a reliable proxy for cellular energy balance (Figure 2F). Surprisingly, the decline in phosphocreatine levels suggested that the energy deficit caused by mitochondrial dysfunction was relatively modest, implying that increased glycolytic activity to high extent compensated for impaired mitochondrial ATP production.

Furthermore, changes in the ratio of SAM/SAH, NAD+/NADH and levels of succinate, cAMP, lactate and other multirole metabolites involved in establishment and control of various epigenetic modifications, sirtuins activity, posttranslational modification and activity of signalling pathways(25–29) clearly indicate that these metabolic changes can have major effect on cellular phenotype (Figure 2G-I). These findings suggest that OPA1 loss can influence macrophage phenotype not only through bioenergetic perturbation but also via metabolic control of gene expression and signaling networks. To delineate polarization-specific metabolic changes induced by OPA1 deficiency (M0 vs. M0, M1 vs. M1, M2 vs. M2) and to compare them with those triggered by classical M1 polarization in control macrophages (M0 vs. M1), we performed pathway enrichment analysis using MetaboAnalyst (Figure 2J). Intriguingly, in OPA1-deficient macrophages, enrichment analysis revealed both the expected mitochondrial-associated pathways such as one-carbon metabolism by folate (linking mitochondrial activity to nucleotide synthesis, amino acid metabolism, and redox balance), fatty acid degradation (FAO), and metabolism of valine, leucine, and isoleucine (branched-chain amino acid degradation) as well as significant alterations in pathways less directly linked to mitochondrial function, including the metabolism of amino acids, purines, pyrimidines, and vitamins.

Notably, several top-enriched pathways were shared between the two comparisons, M0 *OPA1^ΔM^* versus M0 control macrophages and control M1 versus control M0 macrophages, indicating that mitochondrial dysfunction evokes a subset of M1 polarization–associated metabolic programs. Among the most consistently affected pathways were (1) one-carbon pool by folate, (2) alanine, aspartate, and glutamate metabolism, (3) β-alanine metabolism, (8) histidine metabolism, (13) nicotinate and nicotinamide metabolism, (16) pyrimidine metabolism, and (19) arginine biosynthesis. Collectively, these form a metabolic module characteristic of both activation-induced and genetically induced mitochondrial dysfunction.

While our targeted dataset of 173 metabolites provided sufficient coverage to identify major metabolic trends, a comprehensive core metabolome analysis encompassing >500 metabolites will be required to fully resolve the metabolic consequences of mitochondrial dysfunction in macrophages.

### Mitochondrial dysfunction and metabolic remodeling associated with OPA1 loss impair macrophage self-renewal

Given the profound alterations observed in pathways related to global metabolism and nucleotide turnover, we hypothesized that the proliferative capacity of OPA1-deficient macrophages might be disproportionately affected due to bottlenecks in nucleotide biosynthesis and one-carbon metabolism, even in the absence of major energetic deficits. Indeed, the abundance of nearly all nucleotides and their precursors was altered in OPA1-deficient macrophages (Figures 3A, S2A).

**Figure 3.**
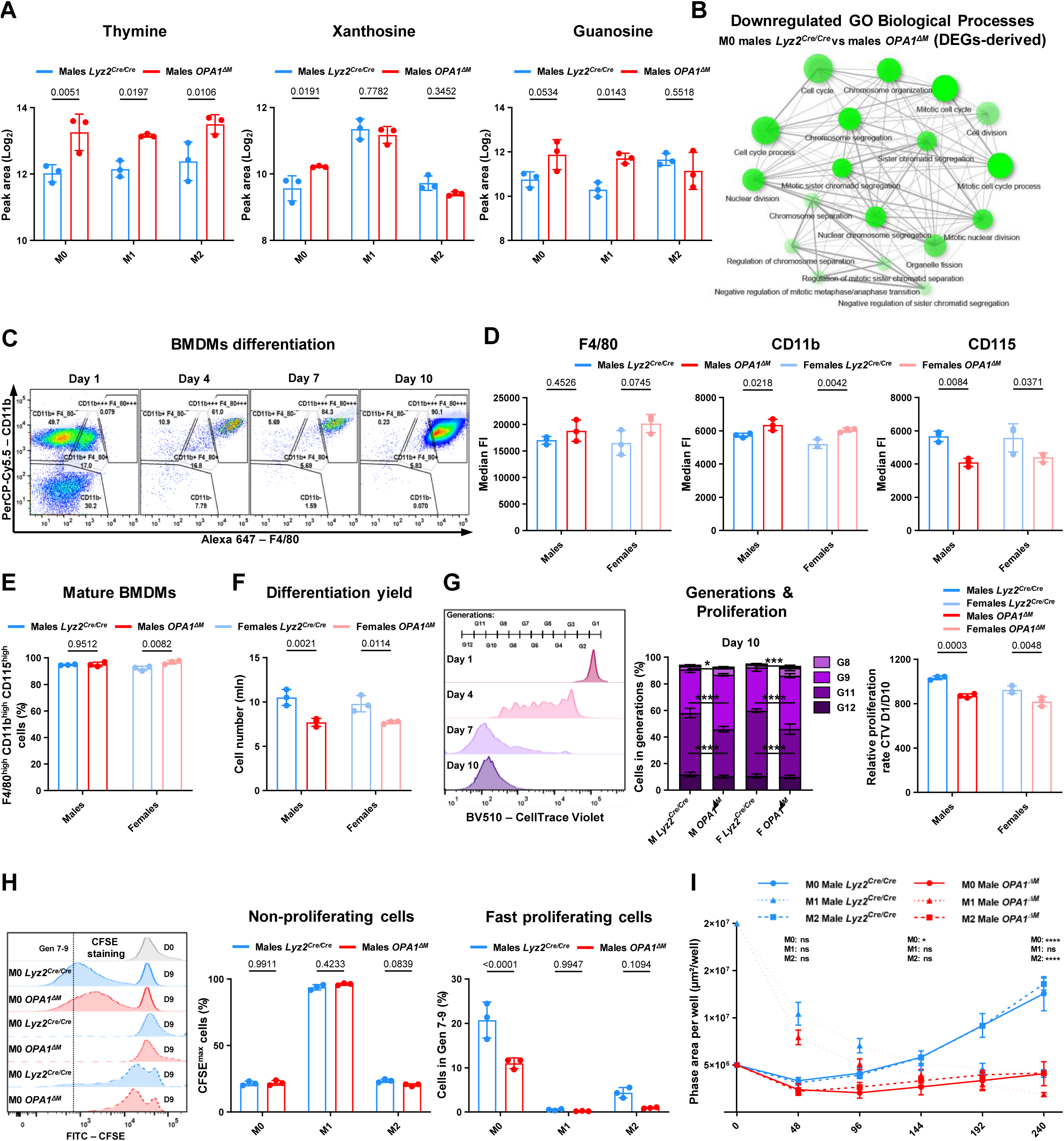
Loss of OPA1 impairs macrophage proliferation. (A) Relative abundances of selected nucleotides and nucleotide precursors identified by targeted metabolomics in polarized male control and *OPA1^ΔM^*macrophages. (B) Enrichment network depicting downregulated biological processes in M0 male *OPA1^ΔM^* macrophages compared with controls, based on differentially expressed genes (RNA-seq). (C) Representative flow cytometry plots illustrating BMDM differentiation stages defined by CD11b and F4/80 expression in control male macrophages. (D) Expression levels of macrophage maturation markers (F4/80, CD11b, CD115) measured by flow cytometry in BMDMs of both sexes and genotypes. (E) Percentage of fully mature macrophages (F4/80^high^ CD11b^high^ CD115^high^) across groups. (F) Total cell numbers indicating macrophage yield at the end of differentiation. (G) Proliferation dynamics of bone marrow progenitors pre-stained with CellTrace Violet; dye dilution was analyzed at days 4, 7, and 10. The relative proliferation rate was calculated as the ratio between the initial and final fluorescence intensity. (H) Fully differentiated BMDMs were stained with CFSE, polarized (M0, M1, M2), and evaluated after 10 days. Fresh medium containing L929-conditioned M-CSF, LPS, or cytokines was replenished every 72 h. The analysis quantified the percentage of non-proliferating cells (retaining CFSE) and fast-proliferating cells (complete dye loss). (I) Long-term male macrophage proliferation monitoring using IncuCyte live-cell imaging over 10 days. For M1 conditions, 1 × 10^5^ cells were seeded; for M0 and M2, 2.5 × 10^4^ cells were seeded. Medium containing L929-conditioned M-CSF, LPS, or cytokines was replenished every 72 h. Data represent mean ± SD; n = 3 (each dot indicates one biological replicate). Statistical significance was determined by two-way ANOVA with Sidak’s correction for multiple comparisons; exact p values are shown within the panels.

To gain mechanistic and functional insights, we analyzed transcriptomic changes by bulk RNA-Seq and enrichment analyses of differentially expressed genes (DEGs). These revealed a pronounced downregulation of gene sets associated with cell cycle progression, strongly suggesting growth inhibition and potential cell cycle arrest.

To verify whether mitochondrial dysfunction affects macrophage differentiation or self-renewal, we next examined the differentiation efficiency of BMDMs from *OPA1^ΔM^* mice. We assessed the expression of canonical macrophage maturation markers (F4/80, CD11b, and CD115) and quantified the proportion of fully differentiated BMDMs in cultures derived from both male and female animals (Figures 3C–E). Our data demonstrated that mitochondrial dysfunction per se does not impair macrophage maturation: OPA1-deficient cells expressed high levels of all canonical macrophage markers (although the expression levels of some markers were altered), and the percentage of F4/80^high^ CD11b^high^ CD115^high^ cells at the end of differentiation was comparable between genotypes. Nevertheless, despite similar marker expression profiles, the overall differentiation yield was substantially reduced in both male and female macrophages derived from *OPA1^ΔM^* mice (Figure 3F), indicating that mitochondrial dysfunction compromises macrophage proliferation during the final stages of differentiation process but without preventing the terminal differentiation itself.

To determine whether the reduced macrophage yield observed in *OPA1^ΔM^*cultures resulted from decreased proliferation, we employed CellTrace Violet (CTV) staining, a fluorescent cell proliferation assay analogous to CFSE. Using this dye, we tracked successive cell divisions (generations) during macrophage differentiation and analyzed proliferation dynamics over time (Figures 3G, S2B).

At day 4 of differentiation, the distribution of cell generations was nearly identical between control and *OPA1^ΔM^* macrophages, indicating comparable proliferation rates at the early stages of differentiation. By day 7, however, initial signs of reduced proliferation became apparent in OPA1-deficient male macrophages, as reflected by a lower proportion of cells in the later generations compared with controls. This effect became more pronounced by day 10, when clear differences emerged in the abundance of cells belonging to the 8th, 9th, and 11th generations, demonstrating that proliferation defects predominantly manifest during the late stages of macrophage differentiation.

Quantification of the relative proliferation rate calculated as the ratio between the CTV signal at the start of the experiment and the residual signal at the final time point confirmed a reduced proliferation rate in *OPA1^ΔM^*macrophages. Consistently, the percentage of cells retaining CTV fluorescence at the end of differentiation was higher in *OPA1^ΔM^* than in control cultures, supporting impaired dilution due to slower or incomplete division cycles.

Importantly, this proliferative defect appeared to be restricted to the late stages of macrophage differentiation, consistent with the view that *Lyz2*-Cre activity is relatively low in circulating monocytes. As a result, OPA1 levels remain comparable between genotypes at the monocyte stage. Accordingly, we did not detect any substantial difference in OPA1 protein abundance using western blotting between monocytes isolated from control and *OPA1^ΔM^*mice by antibody-based magnetic separation (data not shown).

Mature BMDMs are capable of limited proliferation when maintained under M-CSF stimulation in M0 or M2 states; however, upon M1 activation they completely lose this capacity due to cell cycle arrest. To investigate how mitochondrial dysfunction influences the proliferative capacity of mature and polarized macrophages, we pre-stained differentiated macrophages with CTV, subjected them to polarization, and monitored cell divisions over nine days (Figure 3H).

Interestingly, OPA1 loss did not alter the proportion of macrophages capable of entering the cell cycle, indicating that this form of mitochondrial dysfunction alone does not induce complete cell cycle arrest. However, it markedly reduced the proliferation rate: the fraction of cells reaching the latest CTV-diluted generations (no detectable dye) was substantially higher in M0 and M2 control macrophages than in their *OPA1^ΔM^* counterparts.

Consistently, long-term live-cell imaging using the IncuCyte system over 10 days revealed that M0 and M2 control macrophages exhibited progressive increases in cell number, whereas *OPA1^ΔM^* macrophages of both male and female origin displayed a near-complete loss of proliferative expansion (Figure 3I). While M1 macrophages are inherently non-proliferative, OPA1-deficient M1 cells showed a further modest decline in cell number, suggesting that mitochondrial stress associated with OPA1 loss enhances baseline apoptosis.

Collectively, these results demonstrate that mitochondrial dysfunction and the associated metabolic remodeling severely restrict macrophage proliferative capacity. This limitation may be particularly relevant in physiological contexts where macrophage expansion is required, such as during anti-helminth responses or tissue remodeling(30,31). Furthermore, while mitochondrial dysfunction contributes substantially to the impaired proliferation characteristic of M1 macrophages, it alone is insufficient to trigger complete cell cycle arrest.

### Metabolic remodeling induced by OPA1 loss primes macrophages towards M1 phenotype

The effector functions of M1 macrophages are closely linked to the intracellular and extracellular abundance of specific metabolites that can act as signaling molecules to modulate the activity of surrounding cells and shape the inflammatory trajectory. In addition to the increases in lactate and succinate in M0 *OPA1^ΔM^* macrophages (hallmarks of M1 macrophage metabolism that accumulate intracellularly and are released into the extracellular environment) we also detected elevated levels of other M1-associated metabolites, including adenosine, histamine, itaconate, and sialic acid (Figure S3A). The altered abundance of these bioactive metabolites is likely to substantially influence the inflammatory milieu and its resolution dynamics(25,32–36).

The transcription factor NF-κB is a central regulator of M1 macrophage polarization, and its activation state tightly correlates with the gain or loss of pro-inflammatory effector functions(37,38). To assess how mitochondrial dysfunction and the resulting metabolic remodeling affect NF-κB signaling, we evaluated the phosphorylation of NF-κB at Ser536, degradation of its inhibitor IκBα, and the abundance of the NLRP3 inflammasome as an additional indicator of pro-inflammatory activation (Figures 4A, S3B).

**Figure 4.**
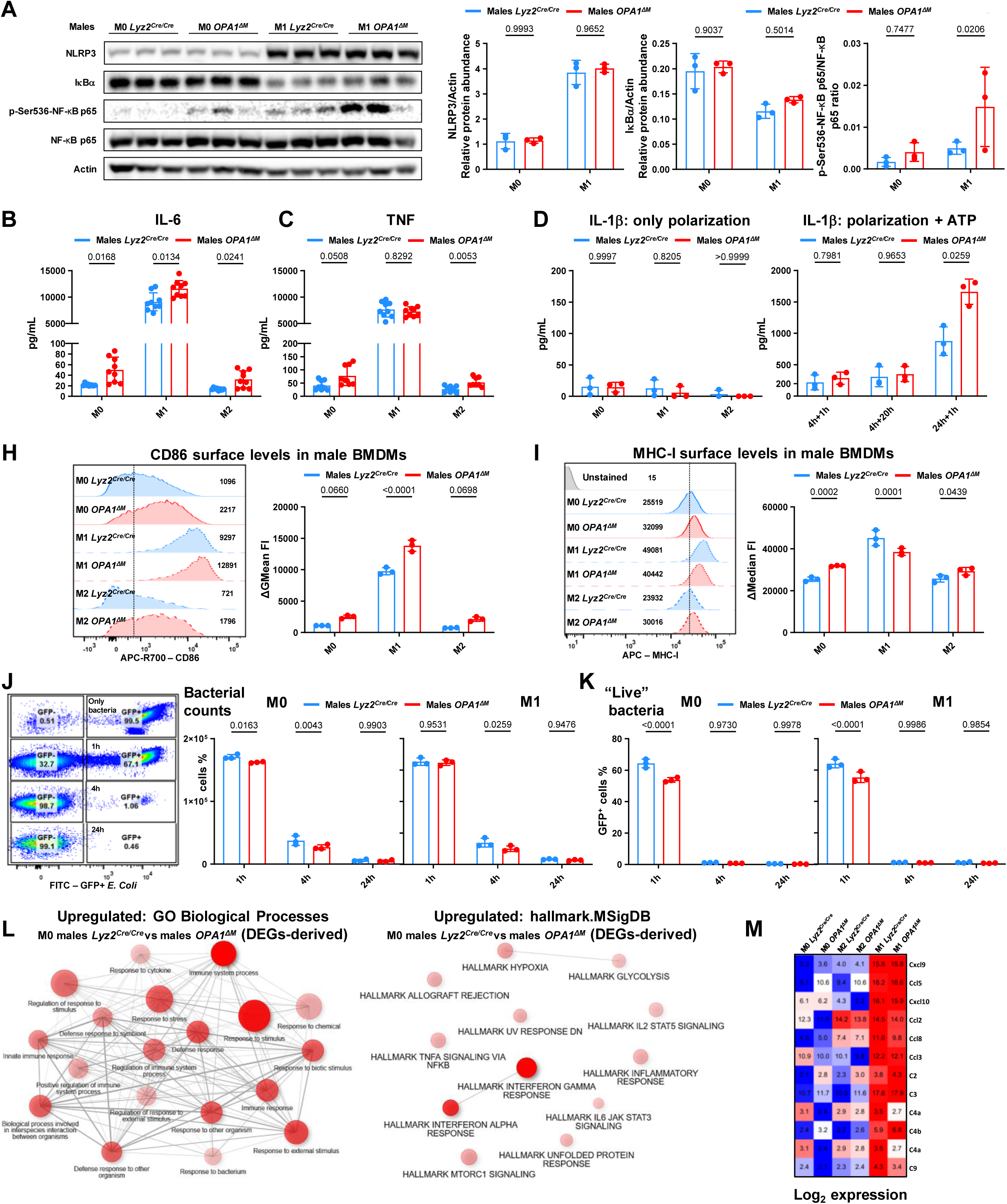
Loss of OPA1 upregulates pro-inflammatory properties in unstimulated macrophages. (A– M) Control and *OPA1^ΔM^* BMDMs differentiated from *Lyz2^Cre/Cre^* and *Opa1^flox/flox^ Lyz2^Cre/Cre^* **male** mice, respectively, were polarized toward M1 and M2 phenotypes or left untreated (M0) and analyzed after 24 h. (A) Western blot analysis of pro-inflammatory proteins in macrophage total lysates. (B, C) ELISA quantification of IL-6 and TNF in supernatants collected from polarized macrophages cultured under high-cell-density, low-volume conditions to enhance autocrine signaling. (D) Left: IL-1β concentrations measured by ELISA in supernatants from polarized macrophages. Right: IL-1β measured after sequential stimulation with LPS + IFN-γ followed by 5 mM ATP for the indicated durations (first value = polarization time; second = ATP exposure). (H, I) Flow-cytometry profiling of M1-associated surface markers on polarized macrophages; cells were pre-blocked with Fc-block. (J, K) Functional bacteria killing assay: M0 and M1 macrophages were co-cultured with GFP^+^ *E. coli* at a 1:50 ratio for the indicated times. (J) Total bacterial counts; (K) frequencies of viable (GFP^+^) bacteria. (L) Enrichment network depicting upregulated biological processes in M0 male *OPA1^ΔM^* macrophages compared with controls, based on RNA-seq differential expression: left – GO Biological Processes library; right – Hallmark MSigDB library. (M) Log₂-normalized expression values showing the mean abundance of selected chemokines and complement components. Data represented as mean ± SD; n = 3–9 (each dot indicates one biological replicate). Statistical significance was determined by two-way ANOVA with Sidak’s correction for multiple comparisons; exact p values are indicated within the panels.

Interestingly, while the abundance of IκBα and NLRP3 remained unchanged, the levels of phosphorylated NF-κB were markedly increased in both male and female *OPA1^ΔM^* macrophages. Whereas NF-κB phosphorylation was barely detectable in M0 control macrophages, clear phospho-NF-κB bands were observed in M0 *OPA1^ΔM^* cells, with particularly strong induction in females. In contrast, among M1 macrophages, NF-κB phosphorylation was more prominently increased in male *OPA1^ΔM^*macrophages, while female M1 macrophages showed minimal differences between genotypes.

To further explore the baseline activation of NF-κB signaling in *OPA1^ΔM^*macrophages, we measured the production of classical pro-inflammatory cytokines whose expression is directly regulated by NF-κB and defines the M1 phenotype, namely IL-6 and TNF (Figures 4C, S3C–D). Strikingly, the abundance of both cytokines was consistently elevated in *OPA1^ΔM^* macrophages under M0 and M2 polarization conditions in both males and females, indicating that the enhanced NF-κB phosphorylation observed in these cells is functionally translated into increased effector output.

Interestingly, while the upregulation of IL-6 and TNF in M0 and M2 macrophages was robust and reproducible across sexes, the effects were much more subtle in M1 macrophages. In M1 cells, IL-6 showed only a minor trend toward upregulation in males and no significant change in females, whereas TNF levels remained unchanged in males and were slightly decreased in female *OPA1^ΔM^*macrophages. Thus, in agreement with our NF-κB activation data, OPA1 loss strongly enhances M1-like inflammatory traits in resting M0 and alternatively activated M2 macrophages but does not necessary further potentiate cytokine production in already polarized M1 macrophages.

We next examined IL-1β, whose production and release are jointly regulated by NF-κB and the NLRP3 inflammasome (Figure 4D, S3E). As expected, IL-1β levels remained near background in all polarization states under basal conditions without ATP stimulation, indicating that mitochondrial dysfunction alone is insufficient to activate the NLRP3 inflammasome. However, upon ATP stimulation, the long-term production and release of IL-1β following M1 polarization were markedly increased in *OPA1^ΔM^* macrophages of both sexes, suggesting that mitochondrial dysfunction primes macrophages for enhanced inflammasome-dependent cytokine responses.

To further confirm that OPA1 loss induces partial acquisition of M1-like features, we analyzed surface expression of M1-associated markers by flow cytometry. Expression of CD86 was significantly upregulated in M0, M1, and M2 macrophages of the *OPA1^ΔM^* genotype, underscoring the strong induction of this marker by mitochondrial dysfunction. In contrast, MHC-I expression followed a pattern similar to cytokine production, an elevation in M0 and M2 macrophages but reduced in M1 macrophages upon OPA1 loss.

Collectively, these findings indicate that moderate mitochondrial dysfunction broadly shifts macrophages towards an M1-like phenotype, enhancing NF-κB activation and pro-inflammatory effector functions in non-activated cells. However, when combined with strong pro-inflammatory stimulation, such as M1 polarization, mitochondrial dysfunction may instead impose excessive stress that limits certain M1-associated activities, shaping a “two-hit” model in which mild mitochondrial dysfunction promotes M1 priming, whereas severe dysfunction can dampen or dysregulates inflammatory outputs.

To assess how the metabolic and inflammatory alterations observed in *OPA1^ΔM^* macrophages translate into functional antimicrobial activity, we exposed control and OPA1-deficient BMDMs either unstimulated or polarized to GFP-labeled *E. coli* and quantified both bacterial phagocytosis and killing (Figure 4J-K, S3F-G).

Surprisingly, the total bacterial burden in co-culture with *OPA1^ΔM^*macrophages was reduced compared with controls. Moreover, the proportion of GFP⁺ bacteria, representing live bacteria, was significantly lower in the presence of OPA1-deficient macrophages. These findings suggest that mitochondrial dysfunction enhances macrophage bactericidal capacity to some extent.

Importantly, this effect could not be attributed to increased phagocytosis. Both the percentage of macrophages actively engulfing bacteria and the intracellular bacterial load per macrophage were reduced in *OPA1^ΔM^*cells, likely reflecting their energy-deficient metabolic state. Thus, the enhanced bacterial clearance observed in OPA1-deficient macrophages appears to result from indirect killing mechanisms, potentially through increased secretion of antimicrobial peptides, reactive intermediates, or other cytotoxic effectors rather than from elevated phagocytic uptake.

Finally, to obtain further evidence supporting the acquisition of pro-inflammatory features in M0 *OPA1^ΔM^* macrophages, we analyzed transcriptionally upregulated processes using our bulk RNA-Seq dataset (Figure 4L). Remarkably, pathway enrichment analysis revealed a broad upregulation of pro-inflammatory gene programs across multiple annotation libraries, indicating robust transcriptomic activation of immune signaling in these cells. Both general immune-related categories such as *immune system process*, *response to cytokine*, *defense response*, *immune response*, *response to bacterium*, and *innate immune response* and specific inflammatory signaling pathways, including *IL-2–STAT5 signaling*, *TNFα signaling via NF-κB*, *interferon-γ response*, *interferon-α response*, and *IL-6–JAK–STAT3 signaling*, were significantly enriched. These results demonstrate a coordinated transcriptional upregulation of innate immune activation pathways in macrophages lacking OPA1.

Moreover, our transcriptomic data revealed altered expression of several chemokines, suggesting potential changes in the ability of OPA1-deficient macrophages to recruit other immune cells. Importantly, we also observed a marked increase in the expression of multiple complement components produced by macrophages, linking the cell-intrinsic activation of these macrophages with broader amplification of humoral and systemic immune responses (Figure 4M).

### Mitochondrial dysfunction mediated by OPA1 loss disrupts M2 macrophage polarization

Because the ability to proliferate, a key determinant of M2 macrophage function during tissue remodeling, was impaired in *OPA1^ΔM^* macrophages, we next investigated how other M2-associated features are affected by mitochondrial dysfunction in OPA1-deficient macrophages.

In contrast to glycolysis-driven M1 macrophages, M2 macrophages are characterized by high OXPHOS, which sustains their long-lived, stable, and metabolically resilient phenotype. This OXPHOS dependence enables M2 macrophages to maintain continuous anabolic and reparative processes, whereas M1 macrophages rely on rapid, glycolytic bursts to execute inflammatory functions at the cost of cell longevity.

Previous studies have highlighted the essential role of FAO in maintaining M2 polarization(39,40). Given the close functional coupling between mitochondria and FAO, it was unsurprising that our metabolomic analysis revealed a marked impairment of this pathway in OPA1-deficient macrophages (Figures 5A–B, S4A). We observed substantial downregulation of L-carnitine, reduced abundance of short-chain fatty acids, and accumulation of long-chain fatty acids, all hallmarks of disrupted FAO and impaired mitochondrial lipid catabolism. These metabolic alterations are consistent with defective M2 polarization.

**Figure 5.**
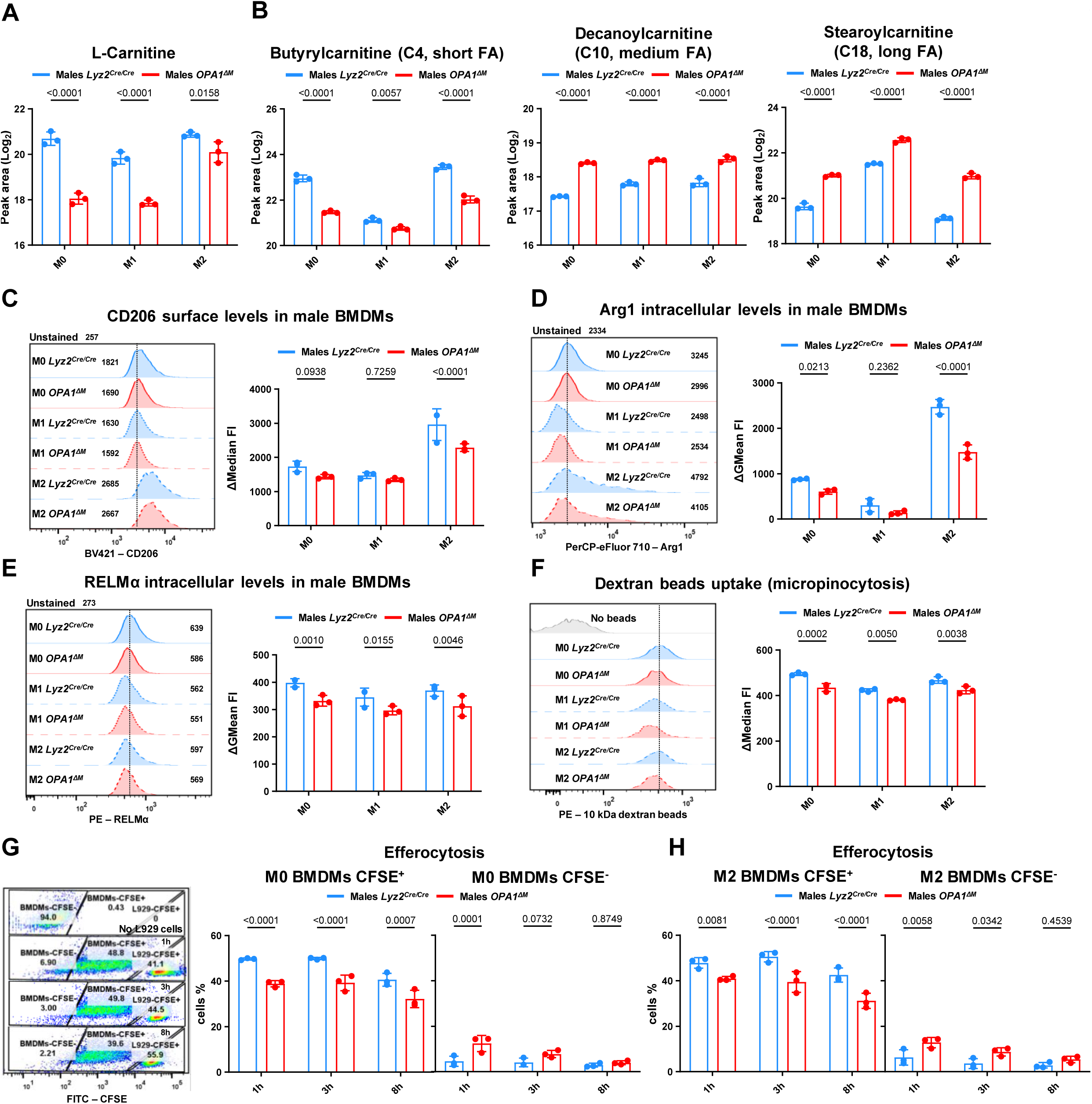
Loss of OPA1 impairs M2 polarization. (A-B) Relative abundances of L-Carnitine and various fatty acids identified by targeted metabolomics in polarized male control and *OPA1^ΔM^*macrophages. (C–H) Control and *OPA1^ΔM^* BMDMs differentiated from *Lyz2^Cre/Cre^* and *Opa1^flox/flox^ Lyz2^Cre/Cre^* **male** mice, respectively, were polarized toward M1 and M2 phenotypes or left untreated (M0) and analyzed after 24 h. (C–E) Flow-cytomety profiling of M2-associated surface markers on polarized macrophages; cells were pre-blocked with Fc-block. (F) Macrophages were incubated with fluorescently labelled 10 kDa dextran–tetramethylrhodamine beads (500 μg/mL) for 30 min, and the fluorescent signal corresponding to bead uptake was quantified by flow cytometry. (G, H) L929 fibroblasts were pre-stained with 5 μM CFSE and rendered apoptotic by incubation at 37 °C for 2 h. Subsequently, apoptotic L929 cells were added to attached M0 or M2 macrophages at a 1:3 ratio for the indicated times. After incubation, wells were washed, trypsinized, and analyzed by flow cytometry for CFSE positivity. Both macrophages (left population) and L929 cells (right population) were detected during acquisition. Data represent mean ± SD; n = 3 (each dot indicates one biological replicate). Statistical significance was determined by two-way ANOVA with Sidak’s correction for multiple comparisons; exact p values are indicated within the panels.

Interestingly, the accumulation and extracellular release of long-chain fatty acids have also been associated with the acquisition of pro-inflammatory properties in macrophages, suggesting that mitochondrial dysfunction may not only hinder M2 maintenance but also actively bias cells toward a more inflammatory state (41,42).

Next, we evaluated the expression of canonical M2 markers such as CD206, Arg1, and RELMα whose abundance closely reflects the extent and stability of M2 polarization. The levels of all three markers were markedly reduced in *OPA1^ΔM^* M2 macrophages in both males and females, indicating impaired acquisition of the M2 phenotype (Figures 5C–E, S4B–D).

We then assessed the efficiency of key “clean-up” functions typically associated with M2 macrophages, including micropinocytosis which we modeled by dextran bead uptake to mimic debris clearance and efferocytosis, the phagocytic removal of apoptotic cells essential for tissue homeostasis. *OPA1^ΔM^* macrophages exhibited a modest but consistent reduction in dextran bead uptake in both sexes, suggesting a mild impairment in macropinocytic activity (Figures 5F, S4E).

For the efferocytosis assay, L929 fibroblasts were pre-stained with CFSE and rendered apoptotic by incubation in PBS at 37°C for 2 hours. In co-culture assays, macrophages efficiently internalized these apoptotic bodies, as reflected by CFSE signal transfer. However, this process was significantly attenuated in both M0 and M2 OPA1-deficient macrophages, as evidenced by a reduced proportion of CFSE⁺ macrophages and a corresponding increase in CFSE⁻ cells (Figures 5G–H, S4F–G).

Collectively, these results demonstrate a consistent downregulation of M2-associated functional programs including proliferation, marker expression, macropinocytosis, and efferocytosis in macrophages with mitochondrial dysfunction, underscoring the central role of mitochondrial integrity in sustaining homeostatic and tissue-remodeling macrophage functions.

### Macrophage mitochondrial dysfunction alters the inflammatory milieu in the peritoneum

To investigate the *in vivo* consequences of macrophage-specific mitochondrial dysfunction, we employed three complementary mouse models that mimic distinct inflammatory contexts and kinetics, and compared them to baseline conditions in control animals. These included: (1) LPS intraperitoneal (i.p.) injection, which induces classical M1-like macrophage polarization; (2) IL-4 complex (IL-4c) i.p. injection, composed of IL-4 and anti–IL-4 antibody, which promotes M2-like polarization and (3) low-grade, age-associated inflammation, representing a chronic physiological inflammatory state(43–45).

Given the sex-specific differences observed in our *in vitro* experiments and the inherent complexity of *in vivo* immune regulation, we analyzed both male and female cohorts across the control, LPS, and aging groups. Using two complementary flow cytometry panels, we profiled immune cell composition in the peritoneal cavity. One panel was designed to assess lymphoid populations, including CD45, CD3, CD4, CD8, CD25, CD69, NK1.1, CD19, CD44, and the neutrophil marker Ly6G, while the second focused on myeloid cells, including F4/80, Ly6C, and CD11c, along with selected polarization markers relevant to macrophage activation states.

PCA and hierarchical clustering of immune cell population frequencies and polarization marker expression revealed that the peritoneal immune landscape was profoundly altered under different inflammatory conditions, with consistent sex-dependent differences (Figures 6A–B). As expected, the LPS model, mimicking acute bacterial inflammation, induced the strongest immunological changes, producing clear segregation of samples and marked separation between male and female mice, underscoring pronounced differences in immune response dynamics between the sexes.

**Figure 6.**
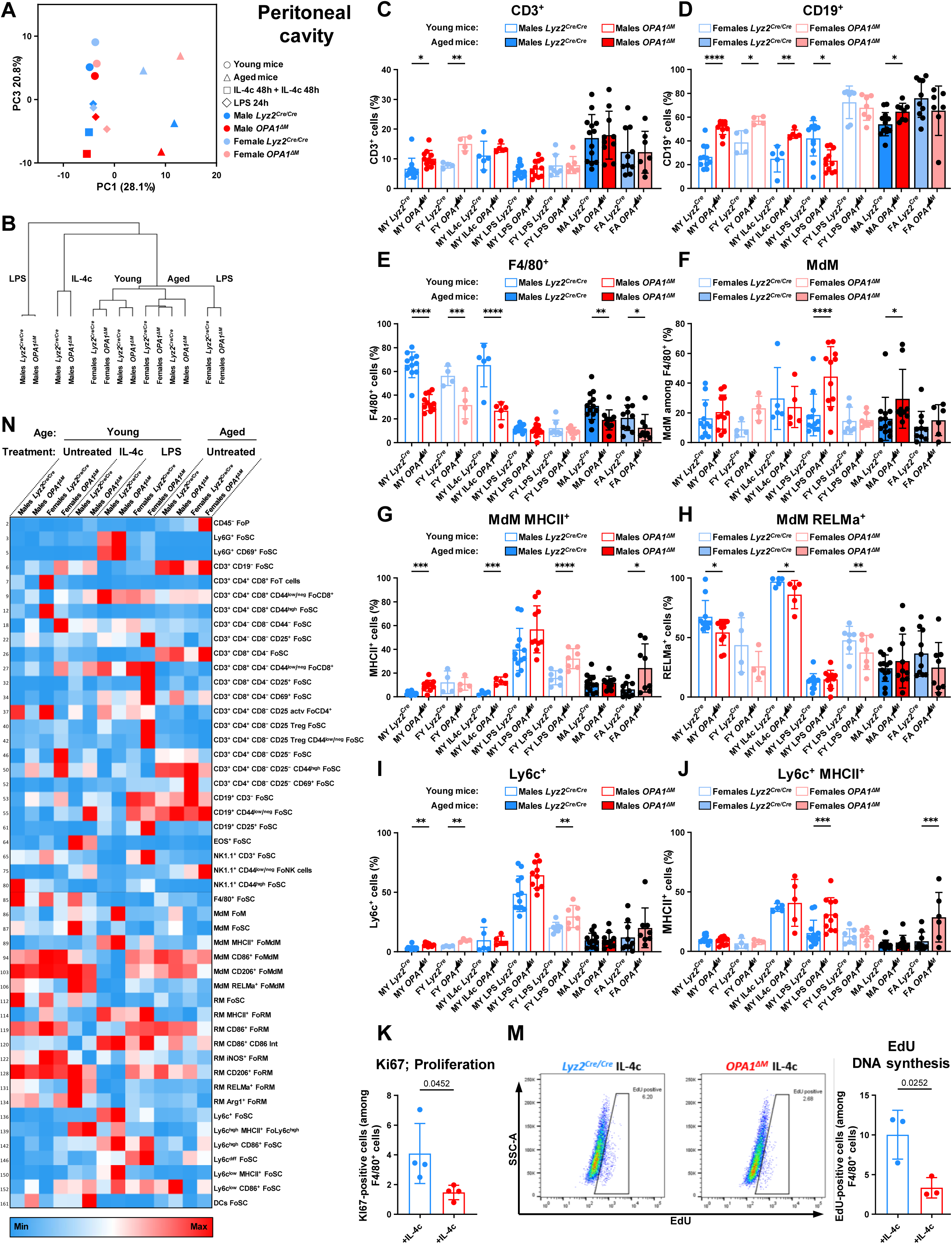
Mitochondrial dysfunction depletes macrophages from the peritoneum. (A-J) Male and female *Lyz2^Cre/Cre^* and *Opa1^flox/flox^ Lyz2^Cre/Cre^*mice were analyzed across seven experimental groups: adult males, adult females, IL-4c–treated adult males (two consecutive intraperitoneal injections of IL-4 complexes; 5 µg recombinant IL-4 complexed to 25 µg IL-4 anti-IL-4 antibody, administered 48 h + 48 h), LPS-treated adult males (10 µg i.p.), LPS-treated adult females (10 µg i.p.), aged males, and aged females. In all groups, peritoneal cells were isolated, Fc-blocked, and profiled by flow cytometry for immune cell composition and macrophage marker expression. (A, B) PCA and hierarchical clustering of immune cell population frequencies and polarization marker expression. (C–J) Frequencies of immune cell subsets positive for the corresponding marker or marker combinations. (K–M) Adult male mice treated with IL-4c complexes (48 h + 48 h) were analyzed for macrophage proliferation using Ki67 staining by flow cytometry or pulsed with EdU (50 mg/kg i.p., 4 h) followed by click-reaction detection of EdU incorporation. (N) Heatmap summarizing representative changes across examined immune populations; each box represents the mean value within a group. Cell populations were quantified relative to parent gates: FoP (frequency of parent), FoSC (single cells), FoT (T cells), FoCD8^+^ (CD8^+^ T cells), FoTCD4^+^ (CD4^+^ T cells), FoNK (NK cells, NK1.1^+^), FoM (all F4/80^+^ macrophages), FoMdM (monocyte-derived macrophages, F4/80^+^ Ly6C^+^), FoRM (resident macrophages, F4/80^+^ Ly6C^-^), and FoLy6C (monocytes, Ly6C^+^ F4/80^-^). Data were collected over multiple years from 29 independent experiments. Values represent mean ± SD; n = 4–13 (each dot indicates one biological replicate); exact p values are shown within the panels.

In young *OPA1^ΔM^* animals, we observed an increased proportion of T and B cells accompanied by a reduction in macrophage abundance, suggesting that mitochondrial dysfunction diminishes the basal macrophage pool even in the absence of inflammatory stimulation, likely due to impaired macrophage self-renewal (Figures 6C–E). In aged animals, this effect was less pronounced, consistent with the overall decline in peritoneal macrophage numbers that accompanies aging.

Given that the peritoneal macrophage compartment consists of two main populations such as tissue-resident macrophages (RM; F4/80⁺ Ly6C⁻) and monocyte-derived macrophages (MdM; F4/80⁺ Ly6C⁺), we next examined their relative contributions (Figure 6F). Our analysis revealed that, in both LPS-induced and age-associated inflammatory settings, male *OPA1^ΔM^* mice displayed a compensatory increase in MdM frequency, likely reflecting an attempt to replenish the macrophage pool in the context of impaired renewal of tissue-resident macrophages.

In line with our *in vitro* results, MdM in *OPA1^ΔM^* mice exhibited increased expression of M1-associated markers, including MHCII (Figure 6G), and reduced expression of M2-associated markers such as RELMα (Figure 6H). However, the magnitude and consistency of these effects varied depending on sex and age.

Consistent with the observed expansion of the MdM population, we also detected enhanced monocyte recruitment to the peritoneum under both basal conditions and following LPS stimulation, though this was not evident in aged animals (Figure 6I). Moreover, these recruited monocytes displayed a more activated phenotype, as indicated by elevated MHCII expression under several examined conditions.

Building on the macrophage-related alterations and our *in vitro* findings, we next assessed how IL-4c treatment, known to drive macrophage proliferation beyond physiological levels *in vivo*, affects the proliferative capacity of OPA1-deficient macrophages. In agreement with the reduced macrophage numbers, Ki67 staining revealed a pronounced decrease in the proportion of proliferating cells in *OPA1^ΔM^* mice. To further quantify DNA synthesis directly, we performed EdU incorporation assays following IL-4c administration. Remarkably, *OPA1^ΔM^* macrophages displayed an approximately 2.5-fold reduction in EdU-positive cells compared with controls, confirming that mitochondrial dysfunction severely limits macrophage self-renewal both at baseline and in response to proliferative immune stimuli.

These deficits appeared to trigger a compensatory increase in monocyte recruitment aimed at replenishing the macrophage pool. A comprehensive summary of the changes in immune cell frequencies and polarization marker expression within the peritoneum is presented in a heatmap overview (Figure 6N).

Collectively, these findings demonstrate that mitochondrial dysfunction in macrophages profoundly alters the peritoneal immune milieu affecting macrophage renewal and polarization and activation, in this way reshaping the broader dynamics of the immune response across multiple cell populations.

### Macrophage mitochondrial dysfunction exerts systemic effects on the immune system

Because blood serves as a central buffer and mirror of physiological status, we hypothesized that macrophage-specific mitochondrial dysfunction could exert systemic effects detectable in circulating immune cells even though macrophages themselves are almost completely absent in the bloodstream. To test this, we applied the same flow cytometry panels used for peritoneal profiling to analyze blood immune composition and activation states.

The overall immune landscape in blood showed clear separation between experimental conditions, comparable to that observed in the peritoneum and these differences were particularly pronounced in challenged animals (LPS-treated or aged) but minimal under baseline conditions, and again showed substantial sex-dependent divergence (Figures 7A–B).

**Figure 7.**
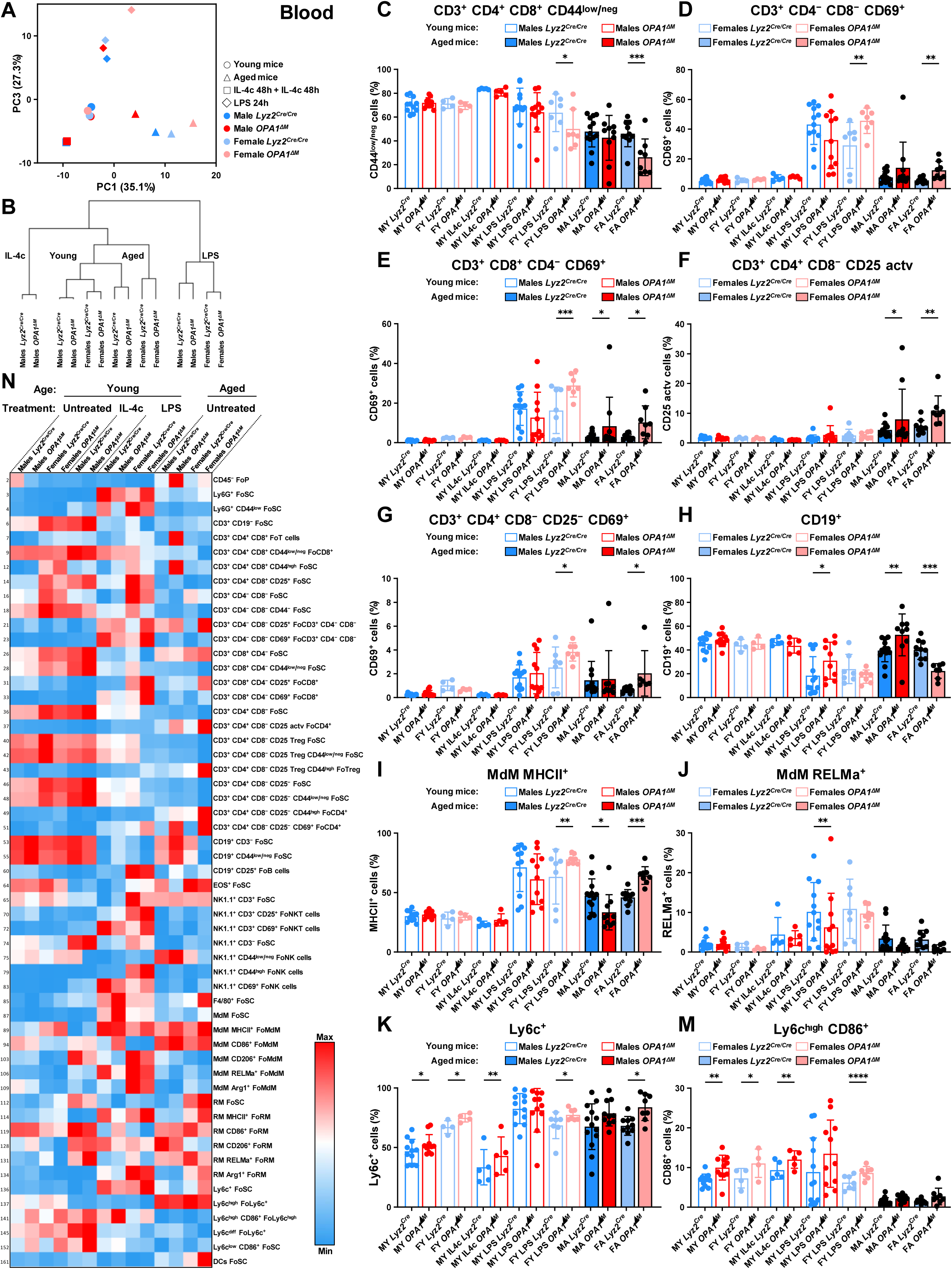
Macrophage mitochondrial dysfunction exerts a systemic effect on the immune system. (A-M) Male and female *Lyz2^Cre/Cre^* and *Opa1^flox/flox^ Lyz2^Cre/Cre^* mice were analyzed across seven experimental groups: adult males, adult females, IL-4c–treated adult males (two consecutive intraperitoneal injections of IL-4 complexes; 5 µg recombinant IL-4 complexed to 25 µg IL-4 anti-IL-4 antibody, administered 48 h + 48 h), LPS-treated adult males (10 µg i.p.), LPS-treated adult females (10 µg i.p.), aged males, and aged females. In all groups, blood cells were isolated, red blood cells were lysed and samples were Fc-blocked, and profiled by flow cytometry for immune cell composition and macrophage marker expression. (A, B) PCA and hierarchical clustering of immune cell population frequencies and polarization marker expression. (C–M) Frequencies of immune cell subsets positive for the corresponding marker or marker combinations. (N) Heatmap summarizing representative changes across examined immune populations; each box represents the mean value within a group. Cell populations were quantified relative to parent gates: FoP (frequency of parent), FoSC (single cells), FoT (T cells), FoCD8^+^ (CD8^+^ T cells), FoTCD4^+^ (CD4^+^ T cells), FoNK (NK cells, NK1.1^+^), FoM (all F4/80^+^ macrophages), FoMdM (monocyte-derived macrophages, F4/80^+^ Ly6C^+^), FoRM (resident macrophages, F4/80^+^ Ly6C^-^), and FoLy6C (monocytes, Ly6C^+^ F4/80^-^). Data were collected over multiple years from 29 independent experiments. Values represent mean ± SD; n = 4–13 (each dot indicates one biological replicate); exact p values are shown within the panels.

Flow cytometry characterization of activation markers in CD3⁺ T cells including conventional CD4⁺CD8⁻ and CD4⁻CD8⁺ subsets, as well as rare double-positive CD4⁺CD8⁺ and double-negative CD4⁻CD8⁻ populations revealed enhanced T cell activation following LPS stimulation and in aged animals, most prominently in female *OPA1^ΔM^* mice (Figures 7C–G). Depending on the subset, these cells displayed a reduced frequency of CD44⁺ T cells, indicating a more chronically activated or memory-like phenotype, together with increased expression of activation markers CD25 and CD69.

Interestingly, B cell dynamics followed an opposite pattern to that observed in T cells. In male *OPA1^ΔM^* mice, the frequency of circulating B cells increased under inflammatory conditions (LPS stimulation or aging), whereas in females, B cell numbers decreased under the same conditions compared to controls. Strikingly, aged animals exhibited a distribution pattern identical to that observed in the peritoneum: in both tissues, macrophage mitochondrial dysfunction increased B cell abundance in males but reduced it in females. These findings suggest that mitochondrial dysfunction in macrophages exerts systemic, sex-specific effects on B cell homeostasis.

Consistent with the enhanced T cell activation observed predominantly in females, the small fraction of macrophage-like cells present in the blood (F4/80⁺Ly6C⁺) also displayed elevated MHCII expression, accompanied by a trend toward reduced expression of the M2-associated marker RELMα (Figures 7I–J).

Furthermore, *OPA1^ΔM^* mice exhibited an increased frequency of circulating monocytes under nearly all examined conditions, suggesting the presence of a compensatory feedback mechanism that enhances monocyte hematopoiesis to replenish MdMs and offset the overall macrophage deficiency (Figure 7K). A substantial fraction of these monocytes also expressed higher levels of CD86, indicating a more pro-inflammatory activation state (Figure 7M).

A comprehensive summary of these systemic immune alterations across all experimental groups is presented in the heatmap (Figure 7N). Notably, most differences in immune composition and activation became apparent only under inflammatory or aged conditions, whereas in the peritoneum some were already detectable at baseline. This observation suggests that macrophage mitochondrial dysfunction exerts cumulative and context-dependent effects that manifest progressively during chronic or systemic inflammation.

## Discussion

Our study, together with two recently published works addressing the consequences of mitochondrial dysfunction in macrophages, establishes a solid foundation for understanding how metabolic reprogramming shapes macrophage immune functions. Our current work, the OPA1-deficient macrophage study by Sánchez-Rodríguez et al. and the Complex I (Ndufs4)–deficient macrophage study by Cai et al. postulate that disrupting mitochondrial function in macrophages elicits a pronounced metabolic reprogramming toward glycolysis (a “Warburg”-like shift). In OPA1-deficient macrophages, both our data and Sánchez-Rodríguez et al. observed a marked reduction in oxidative respiration accompanied by increased glycolytic flux. Similarly, Cai et al. reported that Ndufs4-deficient macrophages exhibit basal oxygen consumption rates and extracellular acidification rates comparable to LPS-activated control cells. This metabolic shift appears to be a general consequence of mitochondrial dysfunction, pushing macrophages toward the bioenergetic state of M1 cells.

Although agreement exists on the metabolic phenotype, the reported immunological consequences diverge. In our hands, OPA1 loss acted as a pro-inflammatory priming signal: *OPA1^ΔM^* macrophages displayed baseline NF-κB activation and elevated secretion of IL-6 and TNF-α in unpolarized and M2 conditions, along with increased CD86, MHC-I, and enhanced IL-1β release upon secondary stimulation. Cai *et al.* described a comparable trend, showing that Ndufs4-deficient macrophages exhibit exaggerated cytokine responses to LPS, consistent with a lower activation threshold. In contrast, Sánchez-Rodríguez *et al.* reported restricted M1 activation owing to reduced IL-6, TNF-α, NOS2/iNOS, and IFNβ expression, suggesting instead a dampened inflammatory potential. These differences likely stem from methodological and biological factors: Sánchez-Rodríguez *et al.* analyzed cells after only 7 days of differentiation and 24 h polarization, whereas we extended BMDM maturation to 10 days in M-CSF–rich conditions (L929) to allow full *Lyz2*-Cre recombination and stabilization of the metabolic phenotype. Early-stage macrophages likely retain residual OPA1 and have not yet undergone complete metabolic and epigenetic remodeling, potentially masking or reversing the effects of mitochondrial dysfunction. Sex-related variability may also contribute, our study identified stronger baseline NF-κB activation in females, whereas sex was not specified by Sánchez-Rodríguez *et al.* and Cai *et al.* noted sex-dimorphic inflammatory responses *in vivo*. Together, these data suggest that mitochondrial dysfunction primes macrophages toward a state of chronic low-grade activation but simultaneously limits the amplitude of acute M1 responses, with context, timing, media composition and sex determining the apparent outcome.

The effects on M2 polarization, however, show clearer agreement between our work and that of Cai *et al.* Both studies demonstrate that mitochondrial dysfunction profoundly compromises IL-4–driven alternative activation. OPA1-deficient and Ndufs4-deficient macrophages alike failed to upregulate canonical M2 markers (Arg1, CD206, RELMα) and exhibited impaired functional processes such as proliferation, debris clearance, and efferocytosis. In our study, *OPA1^ΔM^* macrophages nearly lost their ability to expand in response to IL-4 both *in vitro* and *in vivo*, underscoring their inability to sustain an OXPHOS-dependent reparative phenotype. In contrast, Sánchez-Rodríguez *et al.* reported enhanced expression of M2 markers in OPA1-deficient macrophages, implying an opposite shift. Nonetheless, across both our study and Cai’s, mitochondrial dysfunction consistently emerges as a key constraint on M2 polarization and macrophage self-renewal.

Interestingly, we consistently observed that sex-dependent variation often approached, or even matched, that of the mitochondrial dysfunction itself. When comparing dozens of parameters across macrophages with or without OPA1 loss in both males and females, the average effect size attributable to sex was frequently comparable to that driven by mitochondrial impairment. This indicates that sex constitutes an independent and powerful modifier of macrophage physiology, influencing bioenergetic remodeling, activation thresholds, and downstream immune functions to a considerable degree. Such effect size underscores the necessity of analysing both sexes in mechanistic studies of immunometabolism, as interpretations drawn from single-sex experiments may capture only a subset of the biological variance and risk conflating mitochondrial and sex-specific responses. Interestingly, the gender-related differences in macrophage metabolism also exist in humans: intrinsic metabolic differences between male and female macrophages have been reported in one human macrophage study. According to it, human placental macrophages from female fetuses rely more on FAO at baseline, whereas those from males show higher baseline glycolytic activity (46). Upon LPS challenge, female placental macrophages were relatively resistant to metabolic reprogramming, whereas male macrophages further enhanced their glycolysis.

Lastly the metabolic and immunological features of OPA1-deficient macrophages suggest that mitochondrial dysfunction may enhance organism’s anti-tumour potential. The partial M1-like reprogramming we observed in resting and alternatively activated macrophages likely promotes a more pro-inflammatory and immunostimulatory environment. *In vivo*, these changes were accompanied by a reduced macrophage pool and increased recruitment and activation of T and B lymphocytes. At a tumour site this can imply a systemic shift toward enhanced adaptive immune surveillance and reduced load of tumour associated macrophages. Furthermore, loss of M2 features supporting cancer cells may favor anti-tumour immunity by limiting immunosuppressive macrophage subsets and sustaining inflammatory tone(47). Thus, although mitochondrial dysfunction compromises macrophage longevity, it simultaneously biases the immune system toward a more inflammatory, metabolically alert state that could confer enhanced resistance to tumor growth and progression.

## Materials and Methods

### Mice

*Lyz2^Cre/Cre^ Opa1^flox/flox^* mice were generated and previously described by our group (Amini et al., 2018). Young cohorts in the study were 2–4 months old, while aged cohorts included animals aged 23 and 27 months. Mice were maintained under specific pathogen-free conditions in individually ventilated cages with ad libitum access to food and water, and were regularly monitored for pathogens. All animal experiments were approved by the Cantonal Committee for Animal Experimentation of Bern and conducted in accordance with Swiss animal protection laws.

For acute inflammatory stimulation, mice received 10 µg LPS (Sigma-Aldrich, Cat. # L6529) per 25 g body weight by intraperitoneal injection in 200 µL DPBS, with the injection volume adjusted proportionally to body size. For M2-type stimulation, mice were subjected to double intraperitoneal exposure to IL-4 complexes (IL-4c), each consisting of 5 µg recombinant IL-4 (13.5 kDa; PeproTech) complexed with 25 µg anti-IL-4 antibody (clone 11B11; Bio X Cell), administered 48 h apart in 200 µL DPBS per injection, again adjusted according to animal weight.

### Western blotting

Cells were lysed in buffer containing 50 mM Tris-HCl (pH 7.4), 150 mM NaCl, 10% glycerol, 1% Triton X-100, 2 mM EDTA, 10 mM sodium pyrophosphate, 50 mM NaF, 200 µM Na₃VO₄, 1× PhosSTOP phosphatase inhibitor (Roche, Cat. # 04906845001), protease inhibitor cocktail (Sigma-Aldrich, Cat. # P8340), and 1 mM PMSF. Lysates were incubated on ice for 30 min with intermittent vortexing and sonicated twice for 10 s. After centrifugation (18,000 × g, 15 min, 4 °C), supernatants were collected and protein concentration determined using the Pierce™ BCA Protein Assay Kit (Thermo Fisher Scientific, Cat. # 23225). Proteins (20–50 µg) were denatured in 100 mM DTT and 1× LDS Sample Buffer (Abcam, Cat. # ab119196), separated on SERVAGel TG PRiME gels (SERVA Electrophoresis), and transferred to Immobilon-P PVDF membranes (Merck Millipore, Cat. # IPVH00010). Membranes were blocked with 5% non-fat milk in TBST (0.1% Tween 20, 20 mM Tris, 150 mM NaCl, pH 7.6) for 1 h and incubated overnight at 4 °C with primary antibodies: Phospho-NF-κB p65 (Ser536) (Cell Signaling Technology, Cat. # 3033; rabbit mAb, 1:1000), NF-κB p65 (Cell Signaling Technology, Cat. # 8242; rabbit mAb, 1:1000), p-IκBα (Ser32) (Cell Signaling Technology, Cat. # 2859, clone 14D4; rabbit mAb, 1:1000), IκBα (Cell Signaling Technology, Cat. # 4812, clone 44D4; rabbit mAb, 1:1000), NLRP3/NALP3 (AdipoGen, Cat. # AG-20B-0014-C100; mouse mAb, 1:1000) Pan-Actin (Cytoskeleton, Cat. # AAN01-Apolyclonal, 1:2000) After washing, membranes were incubated with HRP-conjugated secondary antibodies (GE Healthcare Life Sciences) and developed using Immobilon Forte Western HRP substrate (Merck Millipore, Cat. # WBLUF0500) on an Odyssey Fc Imaging System (LI-COR Biosciences).

### Measurement of IL-6, TNF and IL-1β

For ELISA assays, 0.8 × 10^6^ macrophages were seeded per well in 6-well plates and cultured in 1 mL of complete medium. Supernatants collected from macrophage cultures were analyzed for cytokine production. IL-6, TNF and IL-1β concentrations were determined by ELISA using the BioLegend ELISA MAX™ Deluxe Set Mouse IL-6 (Cat. # 431304) and Mouse TNF-α (Cat. # 430904) and IL-1β levels were measured using the BioLegend ELISA MAX™ Deluxe Set Mouse IL-1β (Cat. # 432604).

To assess inflammasome-dependent IL-1β release, macrophages were first primed with LPS and IFN-γ and subsequently stimulated with 5 mM ATP (adenosine 5′-triphosphate disodium salt hydrate; Sigma-Aldrich, Cat. # A2383). All assays were performed according to the manufacturers’ instructions.

### Targeted Metabolomics

Cell pellets were extracted with 80% methanol, sonicated in an ultrasonic bath, and transferred into lysis tubes containing ceramic beads for homogenization using a Precellys Cryolys tissue homogenizer (Bertin Instruments). Lysates were centrifuged (15,000 rpm, 15 min, 4 °C), and the supernatants were evaporated to dryness. Dried extracts were reconstituted in methanol in volumes normalized to total protein content, determined from corresponding cell pellets using the Pierce™ BCA Protein Assay Kit (Thermo Fisher Scientific, Cat. # 23225). Metabolite quantification was performed by ultra-high performance liquid chromatography coupled to tandem mass spectrometry (UHPLC–MS/MS) on an Agilent 6495 iFunnel Triple Quadrupole system operated in Multiple Reaction Monitoring (MRM) mode. Two complementary chromatographic methods (reversed-phase and hydrophilic interaction liquid chromatography) coupled with both positive and negative electrospray ionization were used to maximize coverage. Raw LC–MS/MS data were analyzed using Agilent MassHunter Quantitative Analysis software (version B.07.00). Relative metabolite abundance was calculated from extracted ion chromatogram (EIC) peak areas of monitored MRM transitions. Data quality was evaluated across pooled QC samples analyzed throughout each batch; metabolites with a coefficient of variation (CV) > 30% were excluded from further analysis.

### Assessment of macrophage proliferation and macrophage numbers

Macrophages were washed in DPBS and stained with 5 µM CFSE (Enzo Life Sciences, Cat. # ALX-610-030-M025) in DPBS containing 0.1% FBS for 10 min at room temperature. Cells were then washed twice with DPBS supplemented with 30% FBS and seeded in complete culture medium. Following induction of M1 or M2 polarization, medium containing cytokines or LPS was replenished every two days. After 9 days, cells were detached with trypsin, washed, and analyzed by flow cytometry using a BD FACSLyric™ flow cytometer (BD Biosciences).

For live-cell proliferation monitoring, macrophages were seeded in 96-well plates and imaged using the IncuCyte live-cell imaging system (Sartorius). Cell proliferation was quantified based on the Phase Area per Well (µm²/well) parameter. For M1 conditions, 1 × 10^5^ cells were seeded per well; for M0 and M2 conditions, 2.5 × 10^4^ cells were seeded per well. Medium containing L929-conditioned M-CSF, LPS, or cytokines was replenished every 72 h.

### Analysis of bioenergetic profiles using Seahorse XF96

Macrophages were seeded in 96-well Seahorse XF plates at a density of 4 × 10^4^ cells per well in complete medium. After 4 h, cytokines or LPS were added to induce polarization. Following 24 h of incubation, cells were washed and the medium was replaced with Seahorse XF DMEM (Agilent) containing 12 mM glucose, 2 mM L-glutamine, and 1 mM sodium pyruvate. Plates were then incubated for 1 h at 37 °C in a non-CO₂ incubator to allow temperature and pH equilibration.

Stock solutions of oligomycin, FCCP, rotenone, antimycin A, and 2-deoxy-D-glucose (all from Sigma-Aldrich) were prepared in the same assay medium and loaded into the injection ports of the Seahorse cartridge. Oxygen consumption rate (OCR) and extracellular acidification rate (ECAR) were measured under basal conditions and after sequential injections of 1.5 µM oligomycin, 2 µM FCCP, 1.5 µM rotenone + 1.5 µM antimycin A, and 50 mM 2-deoxy-D-glucose using the Seahorse XFe96 Analyzer (Agilent).

### Flow cytometry analysis of macrophage polarization markers and immune populations in peritoneum and blood

For in vitro experiments, polarized and unstimulated macrophages were washed, trypsinized, washed again, and resuspended in DPBS containing 2% FBS and Fc-blocking buffer (TruStain FcX™ anti-mouse CD16/32; BioLegend). Surface staining was performed for 20 min at 4 °C using the following antibodies: MHC-I–APC (Thermo Fisher Scientific, Cat. # 17-5998-82), CD80–BV605 (BioLegend, Cat. # 104729), CD86–APC-R700 (BD Biosciences, Cat. # 565479), CD206–BV421 (BioLegend, Cat. # 141717). After washing, cells were fixed with 2% paraformaldehyde (Sigma-Aldrich) for 10 min at room temperature, permeabilized with 0.5% saponin in DPBS, and stained intracellularly for 20 min at 4 °C with RELMα–PE (Thermo Fisher Scientific, Cat. # 12-5441-82), iNOS–AF488 (Thermo Fisher Scientific, Cat. # 53-5920-82), and Arg1–PerCP-eFluor 710 (Thermo Fisher Scientific, Cat. # 46-3697-80). After the final washes, samples were acquired on a BD FACSLyric™ flow cytometer (BD Biosciences) and analyzed using FlowJo v10 software (Tree Star).

For in vivo immune profiling, peritoneal lavage and peripheral blood cells were isolated, washed, and Fc-blocked using TruStain FcX™ (anti-mouse CD16/32; BioLegend). Red blood cells in whole-blood samples were lysed using ACK lysis buffer (Thermo Fisher Scientific) prior to staining. Panels were optimized to characterize both myeloid and lymphoid populations as well as activation and polarization markers. The following antibodies were used: Leukocyte and activation markers: CD45– PerCP (BioLegend), CD69–APC (BioLegend, Cat. # 104513), CD25–PE (BioLegend, Cat. # 102007), and MHC-II (I-A/I-E)–BV510 (BioLegend, Cat. # 107635), CD44–BV510 (BioLegend, Cat. # 103044) T-cell and NK-cell markers: CD3ε–APC-Cy7 (BD Biosciences, Cat. # 557596), CD4–V450 (BD Horizon™, Cat. # 560470), NK1.1–KIRAVIA Blue 520™ (BioLegend, Cat. # 56522), CD8-PE/Cy7 (BioLegend) Myeloid markers: F4/80–APC/Cy7 (BioLegend, Cat. # 123117), Ly6C–PE/Cy7 (BioLegend, Cat. # 128017), Ly6G–BV605 (BD Horizon™, Cat. # 563005), CD11c-APC (BioLegend).

### Efferocytosis and bacterial killing assays

Macrophages were seeded in 12-well plates at a density of 2.25 × 10⁵ cells per well in complete medium and allowed to adhere overnight, forming near-confluent monolayers. Following polarization into M0, M1, or M2 states, macrophages were subjected to either efferocytosis or bacterial killing assays as described below.

Efferocytosis assay: To generate apoptotic targets, L929 fibroblasts were pre-stained with 5 µM CFSE (Enzo Life Sciences, Cat. # ALX-610-030-M025) and rendered apoptotic by incubation in PBS at 37 °C for 2 h. CFSE-labeled apoptotic cells were added to adherent macrophages at a 1:3 macrophage-to-target ratio and co-incubated for the indicated durations. After incubation, wells were gently washed three times with PBS to remove non-engulfed apoptotic bodies. Macrophages were then detached with trypsin, washed, and analyzed by flow cytometry (BD FACSLyric™, BD Biosciences). Efferocytosis efficiency was quantified as the percentage of CFSE⁺ macrophages.

Bacterial killing assay: Polarized macrophages were co-cultured with GFP-expressing *E. coli* (MOI = 50:1) for the indicated durations under adherent conditions. After incubation, wells were washed three times with PBS to remove extracellular bacteria. Supernatants containing detached or free bacteria were collected separately and analyzed by flow cytometry to determine bacterial counts and proportion of GFP⁺ (live) bacteria. Macrophages were subsequently detached with trypsin, washed, and acquired under identical flow cytometry settings.

### Measurement of mitochondrial biomass, mitochondrial membrane potential, total ROS, and uptake capacity

Polarized and unstimulated macrophages were washed, trypsinized, washed again, and resuspended in complete medium at a final concentration of 1 × 10⁶ cells per mL.

For assessment of mitochondrial biomass, cells were stained with 200 nM MitoTracker™ Green FM (Thermo Fisher Scientific, Cat. # M7514) for 30 min at 37 °C.

For measurement of MMP, cells were stained with 3 µM JC-1 dye (Thermo Fisher Scientific, Cat. # T3168) for 30 min at 37 °C. The ratio of red (PE channel) to green (FITC channel) fluorescence intensity was calculated to quantify relative MMP.

For evaluation of total ROS, cells were incubated in complete medium containing 5 µM dihydrorhodamine 123 (DHR 123; Thermo Fisher Scientific, Cat. # D23806) for 30 min at 37 °C, followed by immediate acquisition.

For assessment of macropinocytic uptake, macrophages were incubated in complete medium containing 500 µg/mL of 10 kDa Tetramethylrhodamine-labeled dextran beads (Thermo Fisher Scientific, Cat. # D1817) for 30 min at 37 °C.

## Acknowledgments

The authors thank Jacqueline Wyss, Joanna Boros-Majewska, and Corinne Felder for their help with this study.

## Author contributions

N.M. conceived and planned the study, performed experiments, analysed and interpreted data, and wrote the manuscript. A.H., T.F and K.O. executed experiments. A.B., S.Y., C.B., H.U.S. suggested experiments, provided advice, analysed data, shared reagents and edited the manuscript. H.U.S. provided overall guidance, experimental advice, the laboratory infrastructure. All authors read and approved the final manuscript.

## Data and materials availability

All data are available in the main text or the supplementary materials.

---

**Supplementary Figure 1.**
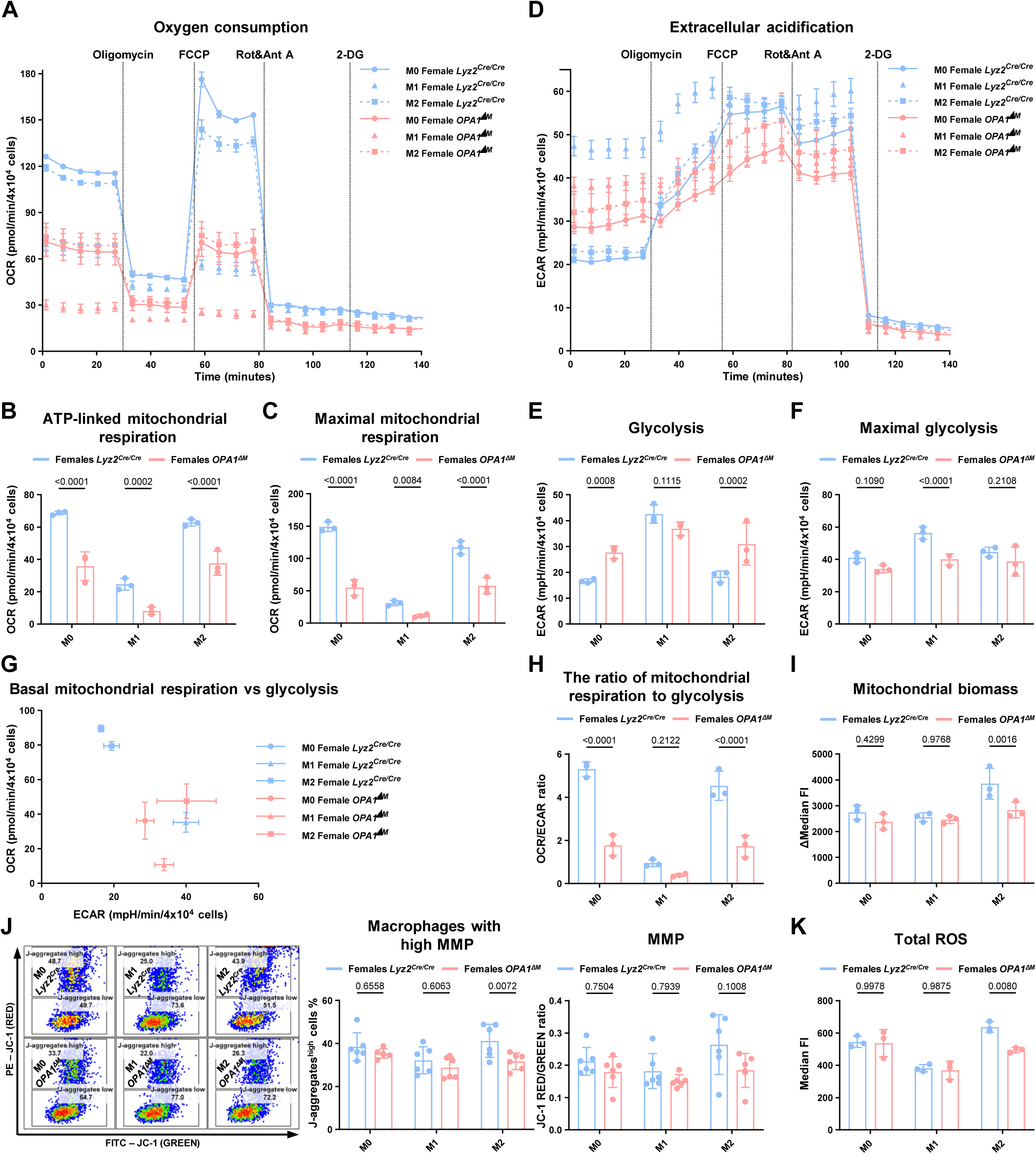
Loss of OPA1 drives a glycolytic shift and reduces mitochondrial respiration in macrophage. (A–K) Control and *OPA1^ΔM^*BMDMs differentiated from *Lyz2^Cre/Cre^* and *Opa1^flox/flox^ Lyz2^Cre/Cre^* **female** mice, respectively, were polarized toward M1 and M2 phenotypes or left untreated (M0) and analyzed after 24 h. (A–H) Bioenergetic profiling using a Seahorse XF analyzer following sequential injections of oligomycin, FCCP, rotenone + antimycin A, and 2-DG. (A–C) Parameters related to oxygen consumption rate (OCR). (D–F) Parameters related to extracellular acidification rate (ECAR). (A, D) Representative Seahorse measurements from one experiment. (B, C, E, F, H) Quantification of OXPHOS- and glycolysis-associated parameters. (I) Mitochondrial biomass determined by flow cytometry using 200 nM MitoTracker Green FM staining. (J) Mitochondrial membrane potential and the proportion of high-MMP cells measured by flow cytometry with 3 μM JC-1 dye. (K) Total ROS quantified by flow cytometry using dihydrorhodamine 123. Data represented as mean ± SD; n = 3–6 (each dot indicates one biological replicate). Statistical significance was determined by two-way ANOVA with Sidak’s correction for multiple comparisons; exact p values are indicated within the panels.

**Supplementary Figure 2.**
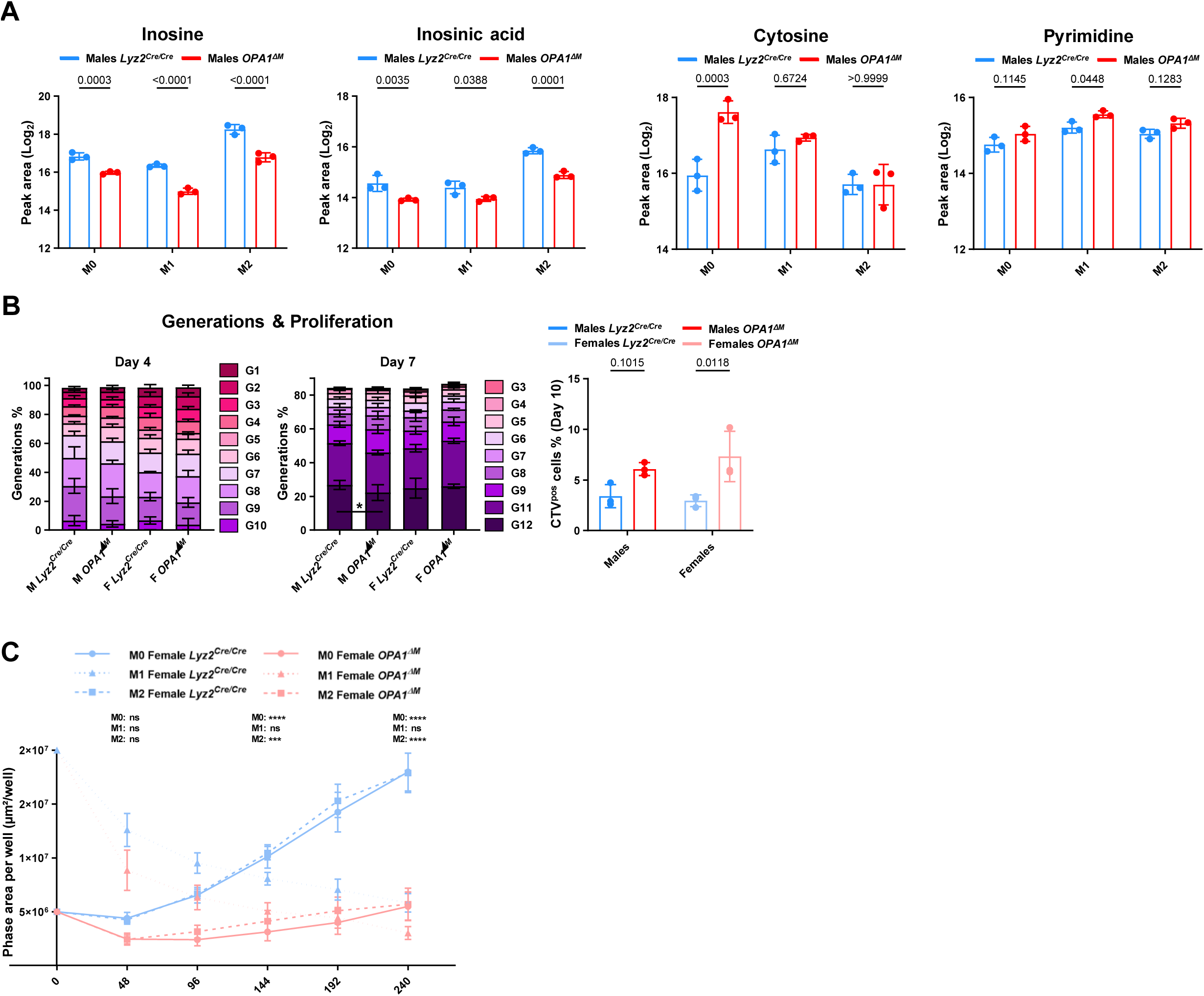
Loss of OPA1 impairs macrophage proliferation. (A) Relative abundances of selected nucleotides and nucleotide precursors identified by targeted metabolomics in polarized male control and *OPA1^ΔM^* macrophages. (B) Proliferation dynamics of bone marrow progenitors pre-stained with CellTrace Violet; dye dilution was analyzed at days 4, 7, and 10. The percentage of dye-positive cells at day 10 reflects the number of slowly-proliferating cells. (C) Long-term female macrophage proliferation monitoring using IncuCyte live-cell imaging over 10 days. For M1 conditions, 1 × 10^5^ cells were seeded; for M0 and M2, 2.5 × 10^4^ cells were seeded. Medium containing L929-conditioned M-CSF, LPS, or cytokines was replenished every 72 h. Data represent mean ± SD; n = 3 (each dot indicates one biological replicate). Statistical significance was determined by two-way ANOVA with Sidak’s correction for multiple comparisons; exact p values are shown within the panels.

**Supplementary Figure 3.**
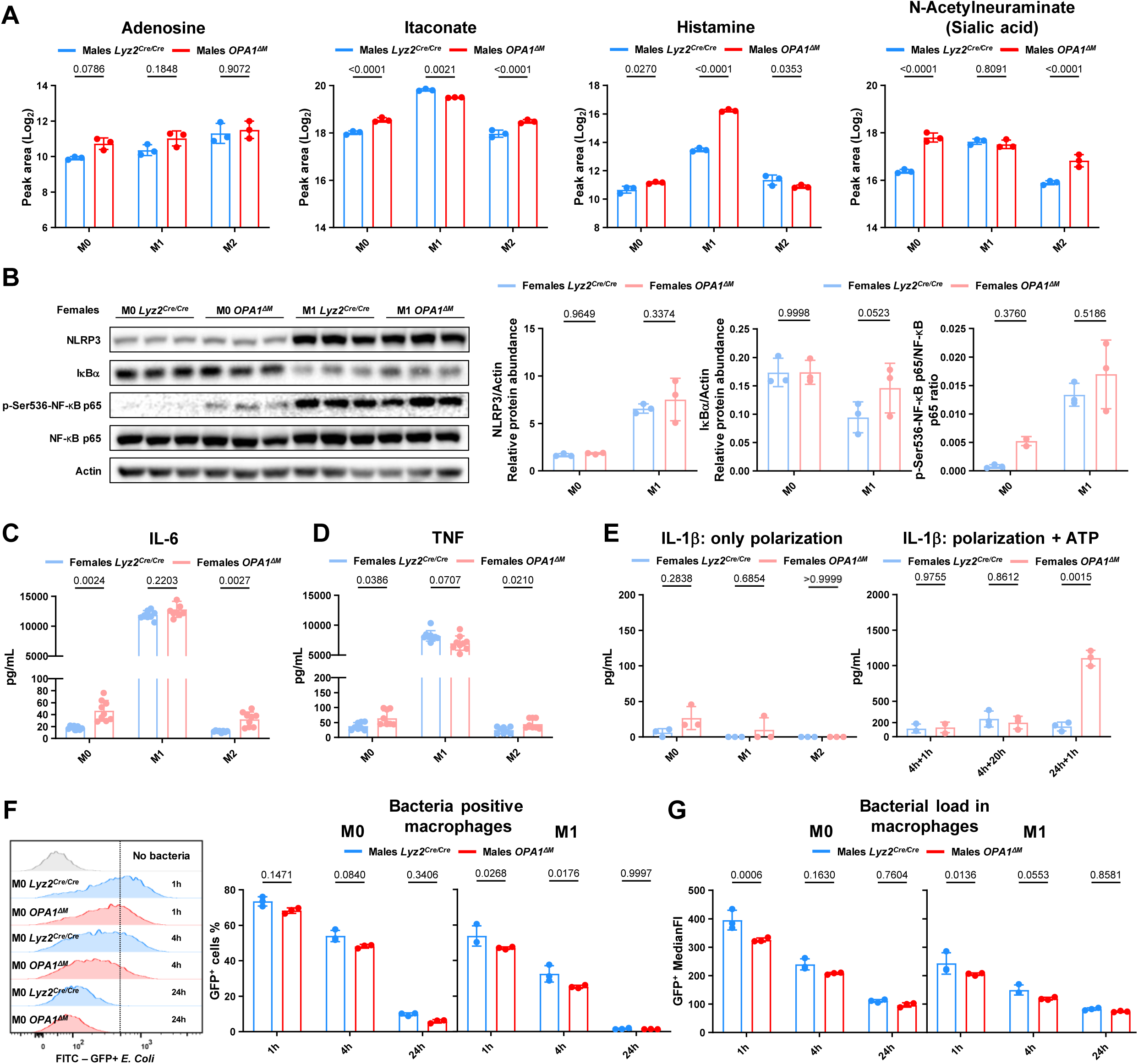
Loss of OPA1 upregulates pro-inflammatory properties in unstimulated macrophages. (A) Relative abundances of inflammation-related metabolites identified by targeted metabolomics in polarized male control and *OPA1^ΔM^* macrophages. (B–E) Control and *OPA1^ΔM^* BMDMs differentiated from *Lyz2^Cre/Cre^*and *Opa1^flox/flox^ Lyz2^Cre/Cre^* **female** mice, respectively, were polarized toward M1 and M2 phenotypes or left untreated (M0) and analyzed after 24 h. (B) Western blot analysis of pro-inflammatory proteins in macrophage total lysates. (C, D) ELISA quantification of IL-6 and TNF in supernatants collected from polarized macrophages cultured under high-cell-density, low-volume conditions to enhance autocrine signaling. (E) Left: IL-1β concentrations measured by ELISA in supernatants from polarized macrophages. Right: IL-1β measured after sequential stimulation with LPS + IFN-γ followed by 5 mM ATP for the indicated durations (first value = polarization time; second = ATP exposure). (F-G) Functional phagocytosis assay: M0 and M1 macrophages were co-cultured with GFP^+^ *E. coli* at a 1:50 ratio for the indicated times. (F) Frequencies of bacteria-positive macrophages; (K) Bacterial “load” represented by GFP signal in macrophages. Data represented as mean ± SD; n = 3– 9 (each dot indicates one biological replicate). Statistical significance was determined by two-way ANOVA with Sidak’s correction for multiple comparisons; exact p values are indicated within the panels.

**Supplementary Figure 4.**
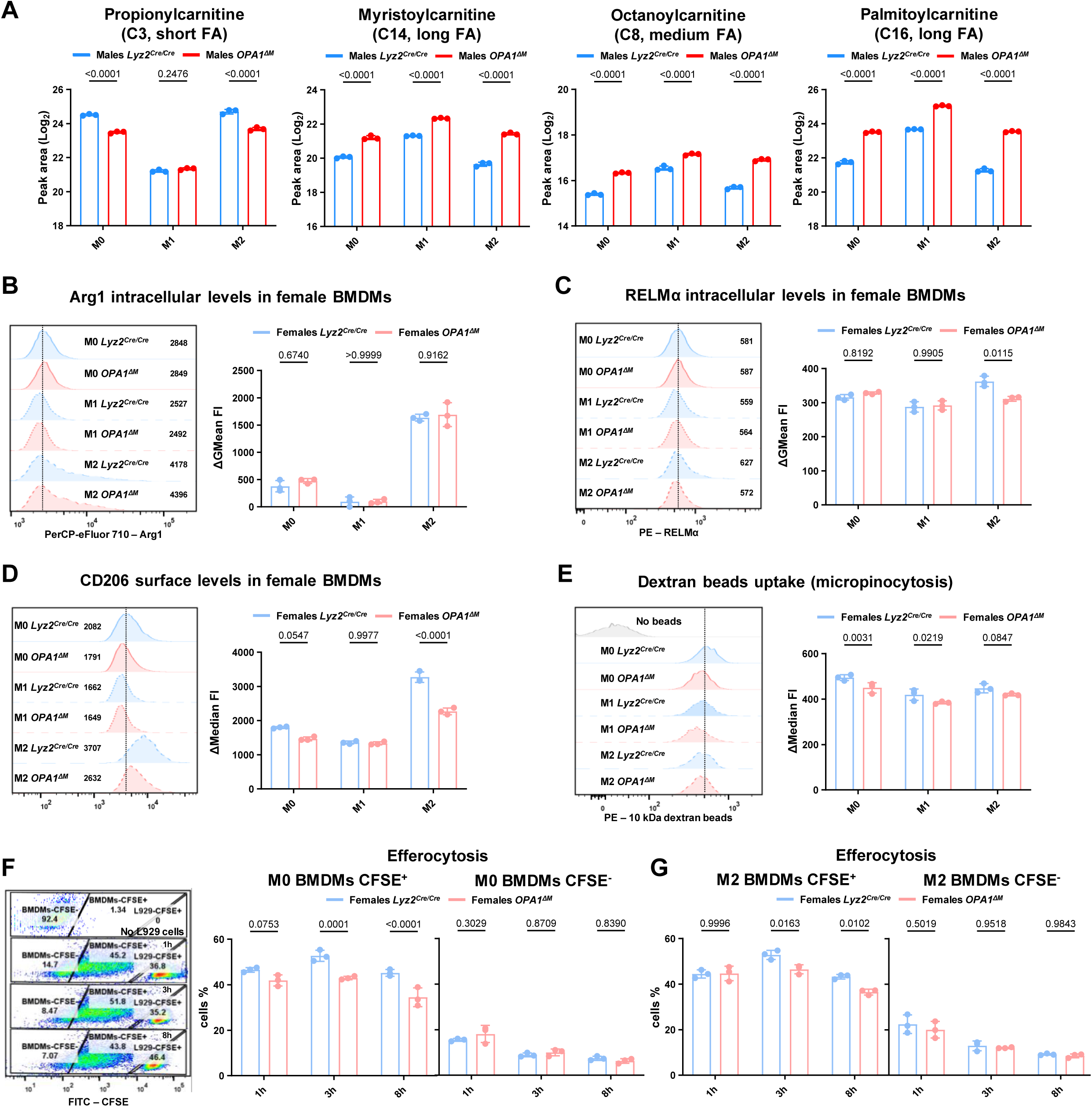
Loss of OPA1 impairs M2 polarization. (A) Relative abundances of various fatty acids identified by targeted metabolomics in polarized male control and *OPA1^ΔM^* macrophages. (B–G) Control and *OPA1^ΔM^* BMDMs differentiated from *Lyz2^Cre/Cre^*and *Opa1^flox/flox^ Lyz2^Cre/Cre^* **female** mice, respectively, were polarized toward M1 and M2 phenotypes or left untreated (M0) and analyzed after 24 h. (B–D) Flow-cytomety profiling of M2-associated surface markers on polarized macrophages; cells were pre-blocked with Fc-block. (E) Macrophages were incubated with fluorescently labelled 10 kDa dextran–tetramethylrhodamine beads (500 μg/mL) for 30 min, and the fluorescent signal corresponding to bead uptake was quantified by flow cytometry. (F, G) L929 fibroblasts were pre-stained with 5 μM CFSE and rendered apoptotic by incubation at 37 °C for 2 h. Subsequently, apoptotic L929 cells were added to attached M0 or M2 macrophages at a 1:3 ratio for the indicated times. After incubation, wells were washed, trypsinized, and analyzed by flow cytometry for CFSE positivity. Both macrophages (left population) and L929 cells (right population) were detected during acquisition. Data represent mean ± SD; n = 3 (each dot indicates one biological replicate). Statistical significance was determined by two-way ANOVA with Sidak’s correction for multiple comparisons; exact p values are indicated within the panels.

